# A Morphodynamic Network Model to Describe Cell Organization and Nematic Ordering in Monolayers

**DOI:** 10.1101/2025.07.21.666048

**Authors:** Prakhar Bandil, Ari Geisler, Claire Leclech, Sara Barrasa-Ramos, Abdul I. Barakat, Franck J. Vernerey

## Abstract

To robustly model nematic ordering in cell monolayers, it is crucial to accurately capture cell geometries, which in turn direct the formation of patterns in cell alignment. In this regard, existing agent-based models (ABMs), such as vertex and Voronoi models, lack generality. Current models often fail to accommodate complex cell geometries (such as elongated or non-convex cells) or implement thermodynamically well-posed rules for cell rearrangements. These features are essential for describing the biophysics of morphogenetic processes where irregular cell shapes and dynamic cell rearrangements play central roles. To address the limitations of existing models, we present a novel computational framework, termed the morphodynamic network model (MNM), for simulating confluent cell monolayers. The MNM is a cell center-based ABM in which the cell centers (nodes) are connected to form a dynamic triangulation (network). The node positions and network structure, recording connections between neighboring cells, evolve according to a split-step approach. First, node positions are updated with the network structure (or topology) fixed, then the network topology is updated with node positions fixed. We apply the MNM to study the morphodynamic behavior of cell monolayers, including fluidization, elasto-visco-plastic responses, nematic ordering, and substrate-induced cell alignment. The MNM is designed to capture complex behaviors and morphologies, offering a flexible and robust tool for studying multicellular phenomena.

## 1. INTRODUCTION

The coordinated movement of cells in dense tissues underlies critical morphogenetic processes such as embryogenesis [1], angiogenesis [2], cancer metastasis [3], and wound healing [4]. Tissue morphogenesis is driven by the dynamics of (1) cell motion and (2) changes in cell count, shape, size, neighbors, and alignment. Recently, numerous studies have demonstrated that cell monolayers exhibit properties characteristic of liquid crystals, such as nematic ordering and topological defects [5–7]. However, in many cases, the mechanisms linking individual cell activity to emergent behaviors at the population scale remain poorly understood. Computational modeling tools can provide unique insights into the spatiotemporal dynamics of cell aggregates. Steered by advancements in computational resources, agent-based models (ABMs) [8, 9], which treat cells as individual agents, have particularly been instrumental. ABMs aim to predict the collective behavior of multicellular systems by defining rules on the agents that govern subcellular (such as actomyosin contractility and cell-cell adhesion), cellular (such as motility, expansion, division, and apoptosis), and multicellular (such as cell rearrangements and collective migration) mechanisms. One of the most extensively used classes of ABMs is the active network model (ANM) [10]. ANMs represent tissue domains as networks of closely packed, deformable, polygonal (polyhedrons in three dimensions) cells. There are two primary archetypes of ANMs: vertex dynamic models (VDMs) [11, 12] and Voronoi tesselation models (VTMs) [13, 14], differing in their degrees of freedom. VDMs employ a network of cell polygon vertices, and links between these vertices define cell boundaries or junctions. In contrast, VTMs define a network (specifically, a Delaunay triangulation) of cell centers. The cell polygons in VTMs are then determined by Voronoi tessellation, a mathematical dual of Delaunay triangulation, of the cell centers. In this case, the cell polygon vertices are given by the circumcenters of the Delaunay triangulation of cell centers. The tissue configuration in ANMs evolves over time through the motion of vertices or cell centers as well as changes in network topology. Over the past decade, several emergent behaviors resulting from local rules have been modeled using these ANMs. Examples include growth [14, 15], flocking [16], viscoelasticity [17], phase transition [13, 18], nematic ordering [19], and other phenomena [20–23].

Despite excelling at capturing some relevant features of morphomechanics, existing ANMs suffer from certain limitations. For instance, VDMs, although highly flexible in terms of cell shapes, do not follow a direct agent-based (or cell-based) framework; instead, they act on subcellular elements, specifically the vertices. Thus, forces from single-cell properties, such as motility [13] and polarity [24], have to be derived from the force balance on the vertices, which are shared by multiple cells [25]. This seems artificial when one seeks to simulate phenomena that arise from individual cell agency, such as collective migration [26]. Additionally, scaling VDMs to large populations (e.g., thousands of cells) or extending them to three dimensions is computationally expensive, as the number of degrees of freedom (vertices) increases significantly with the number of cells. These limitations inherent to a vertex-based approach motivated the development of the cell-center-based VTM (or the self-propelled Voronoi (SPV) model) [13]. However, VTMs require cell polygon configurations to be given by the Voronoi tessellation at all times, imparting strict geometric constraints on cell shape. While VTMs can approximate real cellular configurations in some cases, many geometries, such as elongated and aligned cells, are not well modeled by the Voronoi tiling (as shown in Fig. 1).

**FIG. 1:**
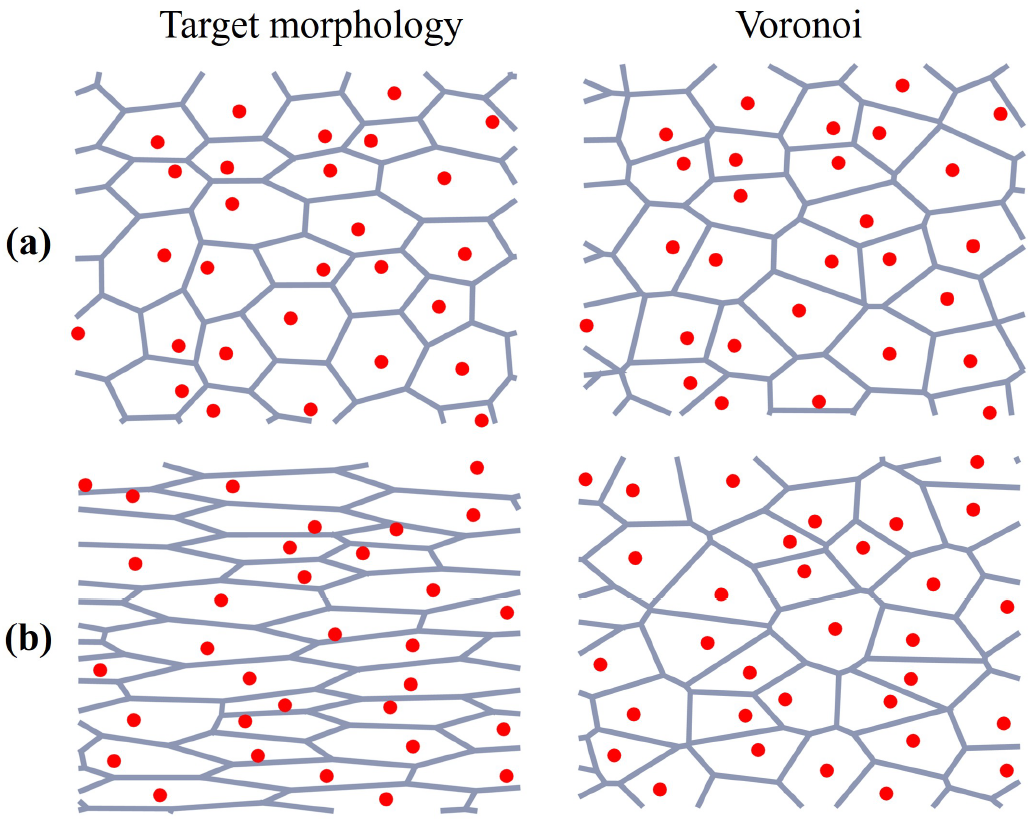
The Voronoi-derived cell tiling for two sets of cell centers with a given target morphology. The VTM does well at capturing roughly cuboidal cells (a), but cannot capture more extreme geometries such as elongated/aligned cells (b).

Existing ANMs also lack an independent timescale for rearrangement dynamics. For instance, in VTMs, the Voronoi tessellation (i.e., the monolayer configuration) is recomputed at every timestep as cell centers move; thus, cells may undergo abrupt rearrangements. These sudden neighbor exchange events, entirely dependent on cell center motion, may not always be desired. Further geometric artifacts resulting from the Voronoi tessellation include rosette formations [14], which are often non-physical. In short, VTMs lack a temporal memory with respect to cell neighbors. In VDMs, cell rearrangements are most frequently modeled by a junction length-based swapping criterion, where if the junction length between two neighboring cells falls below a threshold value, a neighbor exchange event is triggered [11]. Consequently, cell rearrangements in both VDMs and VTMs are entirely dependent on the movement of their respective degrees of freedom. Motion for both models follows over-damped Langevin dynamics with a viscosity coefficient, *η*. Thus, for high *η*, cell rearrangements are suppressed, while for low *η*, cells can rearrange vigorously. As a result, the tissue dynamics depend solely on *η*, which represents all frictional sources. The underlying source of viscous-type friction in a cell aggregate is the breaking and reforming of adhesion bonds. The frictional forces (owing to adhesion) exist at cell-cell junctions as well as at the cell-substrate interface. Thus, there is cell-cell friction (termed tissue viscosity), which we denote by *η*_*cc*_, and cell-substrate friction, which we denote by *η*_*cs*_ [27]. In lumping all friction sources into a single parameter *η*, existing ANMs fail to engage distinct timescales for substrate friction and tissue viscosity, a limitation when modeling systems such as suspended cell monolayers [28], where cell substrate friction is absent altogether, or systems with varying substrate stiffness [27]. The robust modeling of cellular aggregates requires at least two characteristic time scales, one dominated by substrate friction and the other by tissue viscosity.

To further extend upon the insights provided by computational models, the construction of ANMs must be revisited and generalized. This sets the objective for the current study. We propose a novel ANM that follows a cell center-based approach (akin to VTMs), while relaxing the restrictions on cell shape (similar to VDMs) and introducing an independent timescale for cell rearrangements. We refer to the presented model as the morphodynamic network model (MNM). To demonstrate the key features of the MNM, we study two-dimensional cell monolayers; however, the methodology can be readily extended to three-dimensional cell aggregates, such as epithelial shells [29, 30]. The primary goal of the MNM is to accurately capture diverse experimental cell shape distributions, such as shown in Fig. 2, and rearrangement dynamics via a robust framework, enabling a wide-reaching investigation of tissue mechanics driven by cell morphology.

**FIG. 2:**
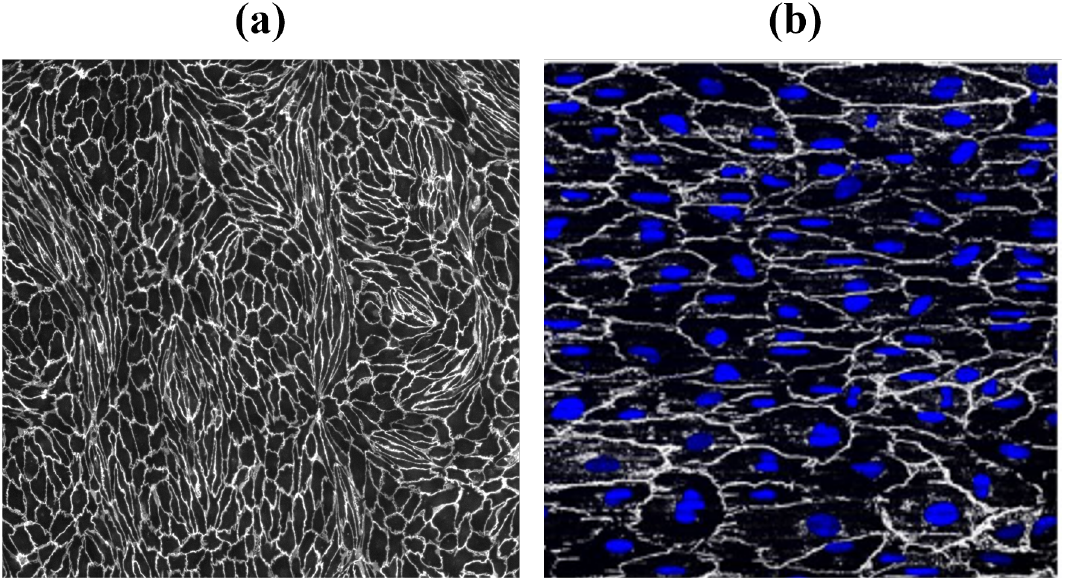
Experimental cell morphologies featuring complex cell geometries not well approximated by the VTM. (a) Elongated endothelial cells after treatment with vascular endothelial growth factor (VEGF); (b) aligned endothelial cells cultured on a microgroove substrate. These systems are investigated further in sections III A and III B, respectively.

The manuscript is organized as follows. In section II, we describe the mathematical framework of the MNM and highlight the model’s properties. We begin by presenting a generalized triangulation approach for constructing polygonal tilings. We then illustrate the dynamics of the monolayer driven by two different modes of cell network reconfiguration. The two modes are: (1) cell center motion, governed by overdamped Langevin dynamics, and (2) topological transitions, modeled using a Metropolis scheme. In section III, we apply the MNM to study some morphodynamic phenomena observed in experiments on endothelial cell monolayers. The results predicted by MNM are found to be in good agreement with the experimental findings. Finally, section IV provides concluding remarks and directions for future work.

## II. MORPHODYNAMIC NETWORK MODEL

### A. Dual network-cell description

As with Voronoi models, the MNM relies on the abstraction of a tissue domain as a network, in particular, a dynamic triangulation of cell centers (or nodes). A dynamic, or temporal, network refers to a network whose structure (defined by its nodes and edges) changes over time. The tissue configuration at any time is given by a polygonal tiling that is a mathematical dual of the underlying triangulation. Suppose that three neighboring cells, *I, J*, and *J* + 1, with centers ***x***_*I*_, ***x***_*J*_, and ***x***_*J*+1_, respectively, form a triangle as shown in Fig. 3. The geometric center of the corresponding triangle then defines the cell polygon vertex, ***r***_*v*_, shared by the three cells. Although various geometric centers, such as circumcenters, incenters, or orthocenters, could be used, we use centroids (defined as ***r***_*v*_ := (***x***_*I*_ + ***x***_*J*_ + ***x***_*J*+1_) */*3), which simplifies cell center force calculations (discussed in section II B 1). Unlike Voronoi models, where the cell center triangulation is always Delaunay, the MNM employs a general triangulation. This means that, unlike Delaunay triangulation, which is unique for a given spatial distribution of cell centers, the triangulation in MNM can assume different structures for the same distribution of cell centers. In doing so, the MNM relaxes the stringent geometric constraints of Voronoi models, enabling the MNM to accommodate a broad range of cell shapes. These include non-convex and highly elongated polygonal cell geometries, as demonstrated later in section III.

**FIG. 3:**
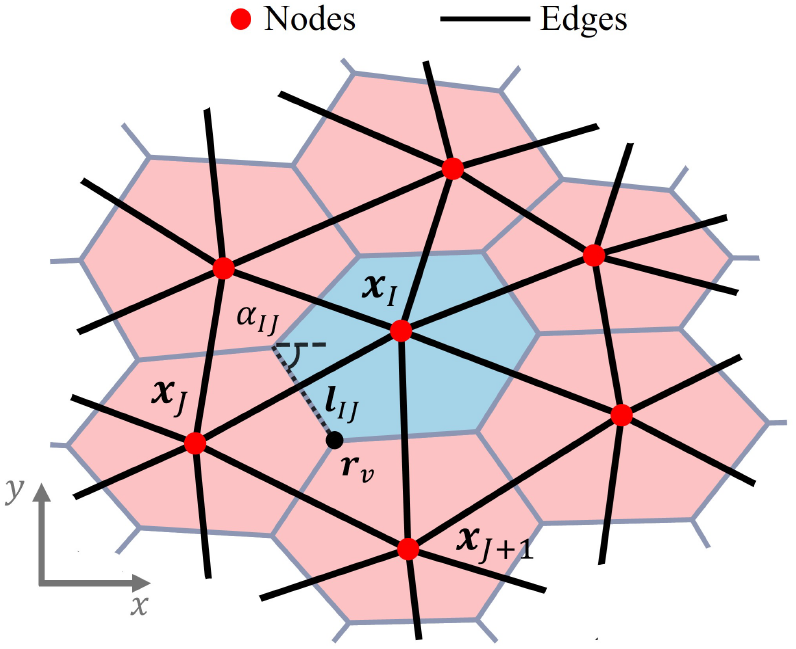
The dual network-cell representation of a monolayer in the MNM, depicting the centroid-derived cell polygon vertex, ***r***_*v*_ (for the triangle of cell centers ***x***_*I*_, ***x***_*J*_, and ***x***_*J*+1_), as well as a junction ***l***_*IJ*_ with length *l*_*IJ*_ and angle *α*_*IJ*_.

In cell center-based ANMs, the triangulation of cell centers conveniently stores information about how cells share junctions, with edges directly corresponding to the connectivity between neighboring (or adjacent) cells. In a dynamic network model, cells can reorganize (i.e., exchange neighbors) by breaking and reforming these edges in a dynamic manner. While this process occurs implicitly in Voronoi models via cell center motion, generalization to non-Delaunay networks requires the MNM to include a novel scheme for updating the network topology. In the MNM, cell neighbor exchange events are modeled by explicitly swapping the edges, independent of cell center motion. Therefore, the cellular configuration in the MNM evolves through two distinct dynamic processes: (1) cell center motion and (2) changes in network topology. To achieve this, MNM employs a decoupled or split-step approach where cell center positions are first updated with the network topology fixed (see Fig. 4(a)), after which the network topology is updated with cell center positions fixed (see Fig. 4(b)). This split-step approach enables us to distinguish the contribution of each process to the morphodynamic response of cell monolayers. In this work, we limit ourselves to non-proliferating tissues (i.e., the number of nodes remains constant). Therefore, the only mode of cell rearrangement considered is the T1 transition, which involves an exchange of neighbors between four cells (as depicted by Fig. 4(b)). We now introduce the MNM in two parts: the first focuses on cell center motion (section II B) and the second on cell rearrangements (section II C).

**FIG. 4:**
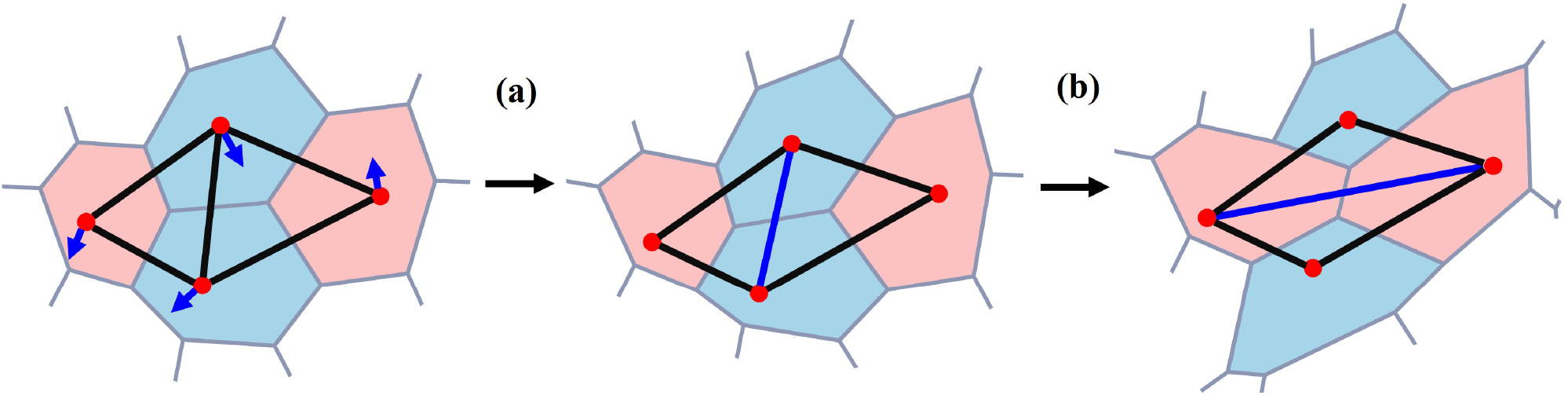
Following four cells through the split-step approach of the MNM, depicting: (a) cell center motion and the resulting changes in cell shape, followed by (b) topological changes given by cell rearrangements (in this case T1 transitions).

### B. Cell center motion

Motion of cell centers within the MNM follows the same governing equations as in the existing ANMs [13, 14]. Therefore, we first provide a brief review of these equations and then explore the elastic response of a monolayer.

#### 1. Updating node position

Consider the position coordinate ***x***_*I*_ (*t*) of the cell center *I* at a time *t*. In the absence of external deformation, the motion of the cell centers is governed by overdamped Langevin dynamics, described by:

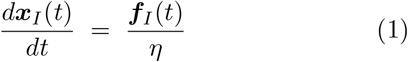

where *d****x***_*I*_*/dt* is the velocity of cell center *I*, ***f***_*I*_ is the total internal force acting on cell center *I*, and *η* is the damping (or viscosity) coefficient. The total internal force ***f***_*I*_ is decomposed into a passive component, 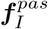, and an active component, 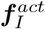, as:

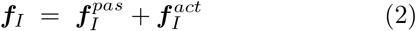

The passive force characterizes the tissue’s equilibrium (or long-term) behavior, while the active component propels the tissue out of equilibrium.

##### Passive force

The passive force is derived from the stored (or elastic) energy *e* as:

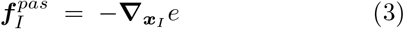

where 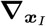 denotes the gradient with respect to cell center position ***x***_*I*_. The passive force acts to minimize the stored energy, driving the monolayer to a stationary ground state. For a polygonal tiling of *N*_*cells*_ cells, the standard energy functional *e* is expressed as [18, 31]:

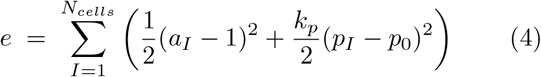

where *a*_*I*_ and *p*_*I*_ are, respectively, the area and perimeter of the polygonal cell. The perimeter modulus *k*_*p*_ models the elasticity of the cell cortex, penalizing changes in cell perimeter. The target shape index *p*_0_ prescribes the preferred shape of cells in the monolayer. Suppose that 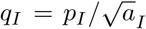 represents the shape index of cell *I*, then the target value of *q*_*I*_ is given by *p*_0_. The utility of *p*_0_ stems from its physical definition [18, 31]:

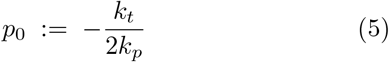

where *k*_*t*_ is the interfacial or junction tension arising from the competition between cell-cell adhesion forces (mediated by E-cadherin junction molecules) and contractile forces (stemming from the actomyosin contractility associated with adherens junctions). While contractile forces work to shrink cell junctions, cell-cell adhesion promotes their expansion. When adhesion dominates, we get *k*_*t*_ < 0 and *p*_0_ *>* 0. Strong adhesion results in higher values of *p*_0_ and larger junction lengths, thereby associating higher values of *p*_0_ with more elongated cell shapes. Situations where contractility dominates (*k*_*t*_ *>* 0 and *p*_0_ < 0) are less common; thus, we focus on *p*_0_ *>* 0.

Equations (4) and (5) assume that all cell junctions are mechanically identical. That is, junction tension *k*_*t*_ is independent of the orientation *α*_*IJ*_ of junction ***l***_*IJ*_ between cells *I* and *J* (see Fig. 3). This assumption is valid in most cases, except for some phenomena (such as convergent extension) that are driven by anisotropic junction tension [32], and modeled using *k*_*t*_(*α*_*IJ*_). We limit ourselves to cases where *k*_*t*_ is isotropic; hence, Eqs. (4) and (5) hold as written.

Having defined the total stored energy *e*, the passive force 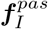 is derived from Eqs. (3) and (4) as [14]:

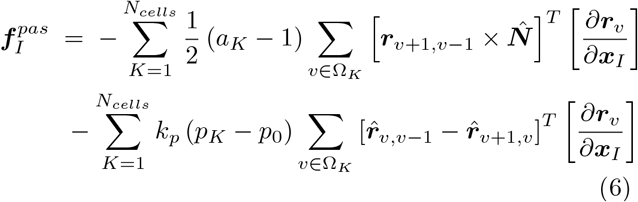

where 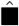 denotes a unit vector in the direction of ■. The vector 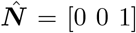 is directed perpendicular to the *xy*-plane of cells. Ω_*K*_ refers to the set of all vertices of cell *K*. The term [*∂****r***_*v*_*/∂****x***_*I*_] is the 3 *×* 3 Jacobian (or transformation) matrix that relates the vertex coordinates to the cell center coordinates.

In this study, the polygonal cell tiling is constructed using the centroids of the cell-center triangles; thus, the Jacobian matrix (for *v ∈* Ω_*I*_) reduces to (1*/*3)***I***, where ***I*** is the 3 *×* 3 identity matrix. For different choices of triangle centers, e.g., circumcenters [14] or orthocenters, the Jacobian matrix must be recalculated accordingly. As depicted in Eq. (6), the total passive force on cell *I* involves contributions from all cells in the monolayer, including cell *I* itself. However, only cell *I* and its immediate neighbors (those sharing a junction with cell *I*) contribute non-zero forces as the Jacobian vanishes for all vertices *v* that do not belong to cell *I* (*v ?∈* Ω_*I*_).

##### Active force

The origin of active forces, 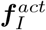, lies in the ability of a cell to migrate with self-propelled velocity (cell motility). The molecular machinery producing active forces is complex; thus, active motion is typically implemented phenomenologically. Considering that the self-propelled speed for cells is *s*_0_, the active force 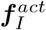 is calculated simply as:

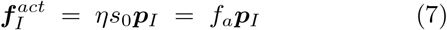

where *f*_*a*_ = *ηs*_0_ is the magnitude of active force and the unit vector ***p***_*I*_ denotes the direction of active cell migration (or cell polarity). Active forces are instrumental in modeling several collective migration phenomena [13, 14]. Cells may migrate along their major principal axis or, similar to Vicsek-type models for active matter, cells could migrate along the average direction of motion of their neighbors. Depending on the mechanism (or rule) implemented for determining ***p***_*I*_, one can explore multiple collective migration phenomena arising from single-cell behavior [14].

##### Updating position

We can now write the MNM’s governing equation for cell center motion. If ***x***_*I*_ (*t*_*n*_) is the position coordinate at time step *t*_*n*_, then the position ***x***_*I*_ (*t*_*n*_ + Δ*t*) at the next time step *t*_*n*_ + Δ*t* is determined by numerically integrating Eq. (1) via a first-order forward Euler scheme as:

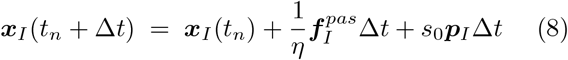

*Note*. (1) All equations presented in this section are formulated in terms of dimensionless quantities. For more details regarding their units, refer to Appendix A. (2) Throughout the study, all monolayer simulations were performed with periodic boundary conditions and assuming no translational activity (i.e., *f*_*a*_ = 0).

#### 2. Elastic response of suspended monolayer

Epithelial tissues, which form the lining of both our internal and external organs, are routinely subjected to large deformations. Notably, epithelial tissues exhibit a complex elasto-visco-plastic response. Over short time scales (typically on the order of minutes), their behavior is purely elastic, with no topological rearrangements (which generally occur over timescales ranging from hours to days [33]). Experiments performed in the elastic regime involve stretching of suspended monolayers and characterizing the stress-strain behavior [34]. We now explore the elastic response of suspended monolayers under stretching as predicted by the MNM.

##### Mechanical stress

We start by defining the mechanical state of stress ***σ***_*I*_ for cell *I*, given by the Batchelor stress tensor as [35]:

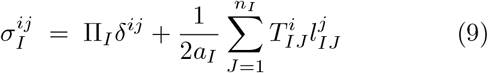

where *i* and *j* denote indices, and the summation in the second term is over all the neighbors *n*_*I*_ of cell *I*. The pressure Π_*I*_, calculated as,

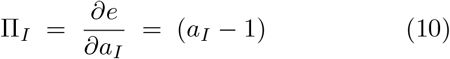

arises from the difference between the cell area and its preferred value. Π_*I*_ < 0 denotes compression and Π_*I*_ *>* 0 denotes tension. 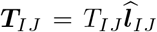 is the tension vector in the junction (or interface) between cell *I* and its neighbor *J*, with:

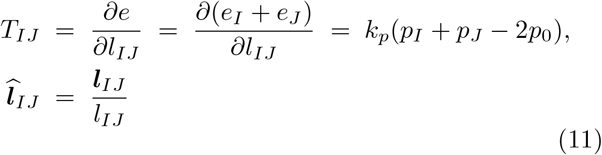

noting only *e*_*I*_ and *e*_*J*_ depend on the junction length *l*_*IJ*_. The global (average) stress ***σ*** in the tissue is then given by:

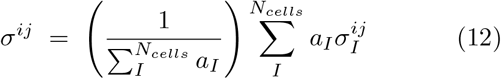

##### Ground state

To investigate the mechanical response of cell monolayers undergoing large deformation, we must establish the ground state of the tissue. The ground state acts as the reference configuration for measuring deformation. For this example, the ground state is taken to be a regular hexagonal lattice (crystalline phase) of cells (see Fig. 5(a)). In this situation, we have perimeter *p*_*I*_ = *p*, area 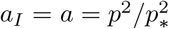, and target shape index *p*_0_ < *p*_*\**_, where 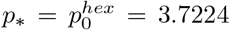 is the shape index for a regular hexagon. Mechanically, the ground state is characterized by ***σ***(*t*_*n*_ = 0) = **0**, which for a crystalline monolayer yields the following cubic equation in perimeter *p* (derivation in Appendix C):

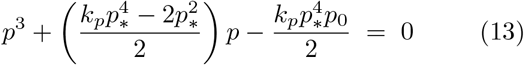

**FIG. 5:**
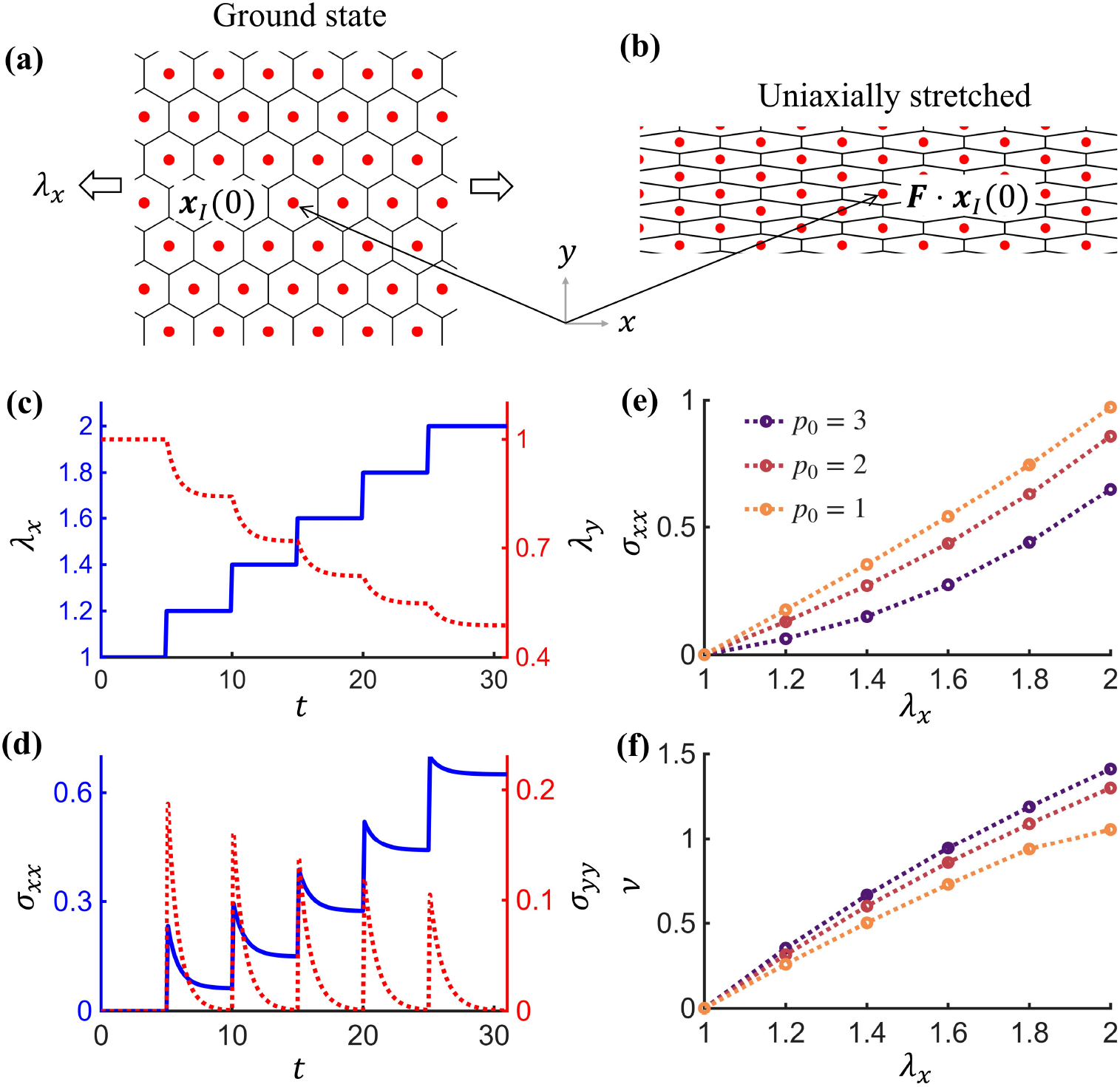
The elastic response of a crystalline monolayer under uniaxial tension, showing: (a) ground state; (b) stretched state; (c) longitudinal stretch *λ*_*x*_ (in solid blue) versus time *t* is applied as a step loading. Lateral stretch *λ*_*y*_ (in dotted red), in response to barostatting (*σ*_*yy*_ = 0), versus time *t* is plotted on the same graph (for *p*_0_ = 3); (d) Stress components, *σ*_*xx*_ and *σ*_*yy*_, versus time *t* (for *p*_0_ = 3). (e) Longitudinal stress *σ*_*xx*_ versus longitudinal stretch *λ*_*x*_ for different values *p*_0_; (f) Poisson’s ratio *ν* versus longitudinal stretch *λ*_*x*_ for different values of *p*_0_. Other simulation parameters: *N*_*cells*_ = 576, *k*_*p*_ = 0.1, Δ*t* = 0.001.

The solution 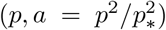 of Eq. 13 for monolayer properties *k*_*p*_ and *p*_0_ is the ground state of the crystalline monolayer. For this problem, we chose *p*_0_ = 3 and *k*_*p*_ = 0.1 (following [15, 31, 36]), resulting in *p* = 3.5251 *> p*_0_ and 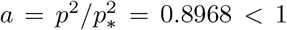 for the crystalline ground state. Substituting these values into Eq. (4), we see that the ground state is not given by the absolute energy minimum *e* = 0, even though ***σ***_*I*_ = **0**. Mechanically, this state is deemed incompatible as cells cannot simultaneously satisfy *e*_*I*_ = 0 and ***σ***_*I*_ = 0. Instead, mechanical equilibrium (***σ***_*I*_ = **0**) is achieved via a complex intracellular stress distribution, wherein the tension forces *T*_*IJ*_ in the junctions (owing to *p > p*_0_) are balanced by the compressive stresses Π_*I*_ inside the cells (owing to *a* < 1) [31, 36, 37]. Having determined the ground state for a crystalline tissue, we can now apply external deformation to a suspended monolayer. Mimicking experimental conditions [34], we perform a uniaxial tensile test on a crystalline tissue using the MNM.

##### Uniaxial tensile test

Let the deformation gradient ***F*** (*t*) at any time *t* = *t*_*n*_ *×* Δ*t* be given by ***F*** (*t*) = *{*diag *λ*_*x*_(*t*), *λ*_*y*_(*t*) *}*, where *λ*_*x*_ and *λ*_*y*_ are stretch ratios of the simulation box along the *x* and *y* directions. In the presence of external deformation ***F***, the motion of cell centers is governed by both the deformation gradient ***F*** and internal forces ***f***_*I*_. Under the deformation gradient, cell centers follow an affine motion given by:

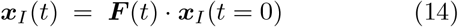

When ***F*** is diagonal (such as in uniaxial stretching), cells initially in the crystalline state maintain their symmetric geometries while becoming elongated under the affine motion of cell centers (see Figs. 5(a) and 5(b)). It can be shown that owing to this symmetry, the passive forces vanish (i.e., 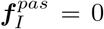). As a result, the motion of cell centers is entirely governed by Eq. (14).

The loading conditions in the uniaxial tensile test are displacement-controlled along the *x* direction and traction-free along the *y* direction. Therefore, while the longitudinal stretch *λ*_*x*_(*t*) is known, the lateral stretch *λ*_*y*_(*t*) needs to be determined from the condition *σ*_*yy*_ = 0, where *σ*_*yy*_ is the lateral component of global stress tensor ***σ***. To implement traction boundary conditions in discrete simulations, a barostat is commonly used. Suppose the stretch, *λ*_*x*_(*t*), is held constant at a specific value. The barostat will then relax the monolayer by deforming the *y* boundary to attain *σ*_*yy*_ = 0 (refer to Supplementary Information (SI), section 1 for more details regarding barostatting). This ‘relaxed’ or the ‘steady-state’ configuration at the given stretch *λ*_*x*_ is then the actual elastically deformed state of the monolayer. To allow sufficient time for the barostat to act, the longitudinal stretch *λ*_*x*_ is applied as a step load as shown in Fig. 5(c). The material properties first considered are *k*_*p*_ = 0.1 and *p*_0_ = 3. We then plot *σ*_*yy*_ versus time *t* in Fig. 5(d), noting that *σ*_*yy*_ completely relaxes to 0 between each successive load step. The corresponding lateral stretch *λ*_*y*_ is also plotted (see Fig. 5(c)). The (actual) stress state for the elastic monolayer under uniaxial stretch is then given by the steady-state values of the longitudinal stress component *σ*_*xx*_ (see Fig. 5(d)).

Using the steady-state values of stress *σ*_*xx*_ corresponding to stretch ratios *λ*_*x*_ during the elastic loading, we plot the stress-stretch curve in Fig. 5(e) characterizing the elastic response of the suspended monolayer. In (qualitative) corroboration with experiments, the MNM predicts a strain-hardening response, where the slope of the curve (denoting stiffness) increases with *λ*_*x*_. We also investigated the effect of target shape index *p*_0_ (keeping *k*_*p*_ fixed at 0.1) on the elastic response of crystalline tissue. The ground states corresponding to properties *k*_*p*_ = 0.1, *p*_0_ = 2 and *k*_*p*_ = 0.1, *p*_0_ = 1 are given by *p* = 3.20, *a* = 0.74 and *p* = 2.7772, *a* = 0.5566, respectively. We observed that the elastic response is stiffer for lower values of *p*_0_. Smaller values of *p*_0_ correspond to weaker cell-cell adhesion. Therefore, the results are counterintuitive as they predict that tissue stiffness increases with a decrease in adhesion. This behavior can be attributed to the anomalous role of adhesion in biological materials, in comparison to passive materials [18, 38]. Another plausible explanation may lie in the complex distribution of subcellular stress. Although the ground state for all three cases (*p*_0_ = 1, 2, 3) is the same (i.e., ***σ***_*I*_ = **0**), the subcellular stress distributions are different with (*p*_0_ = 1, *a* = 0.5566) cells under larger compression than (*p*_0_ = 3, *a* = 0.8968) cells. As a result, the overall elastic behavior could depend on how these complex stress fields interact with the applied external deformation.

We also plot, in Fig. 5(f), the Poisson’s ratio *ν* (calculated as *ν* = − ln(*λ*_*y*_*/λ*_*x*_)) versus stretch ratio *λ*_*x*_. We observe that Poisson’s ratio decreases with a decrease in target shape index *p*_0_. Further decreasing *p*_0_ to extremely low values results in (not shown here) negative Poisson’s ratios (auxetic material). Such a regime has also been reported in both the vertex [39] and continuum [37] models, further validating the elastic response captured here using the MNM. For completeness, we also explored the role of non-affine motion on the elastic response of tissues. For this, we deform a tissue in its amorphous state, resulting in non-zero internal forces. Note that motion under external deformation ***F*** is still affine, but the overall motion would be non-affine due to internal forces. The methodology and simulation results are presented in detail in section 2 of the SI. In brief, we observed that non-affine motion yields a softer elastic response (compared to purely affine motion in the crystalline case), as is commonly observed [36]. This softer response (reduced stress for the same stretch) is attributed to energy dissipation resulting from the motion of cells under (passive) internal forces toward lower energy states.

### C. Cell rearrangements: T1 transitions

So far, we have examined elastic networks that maintain their topology over time. The next step is to investigate the behavior of networks that can undergo topological changes via T1 transitions. We now elucidate how T1 transitions are modeled within the MNM framework.

#### 1. Updating topology

Updating topology in the MNM refers to obtaining a new triangulation for the same spatial distribution of cell centers. The rule for T1 transitions within the MNM follows the Metropolis algorithm, an archetype of Markov chain Monte Carlo (MCMC) methods. The same algorithm is used in the well-known cellular Potts model (CPM) [40, 41]. In the MNM, a monolayer with a given fixed spatial cell center distribution, 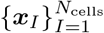, can assume different topological states, say *γ*, corresponding to different triangulations, *𝒯* (see Figs. 4(b) and 6). MCMC is a robust computational tool that enables the MNM to explore the highly complex probability space of network states (in this case, monolayer morphologies/triangulations) for a distribution of cell centers. The scheme is demonstrated as follows.

**FIG. 6:**
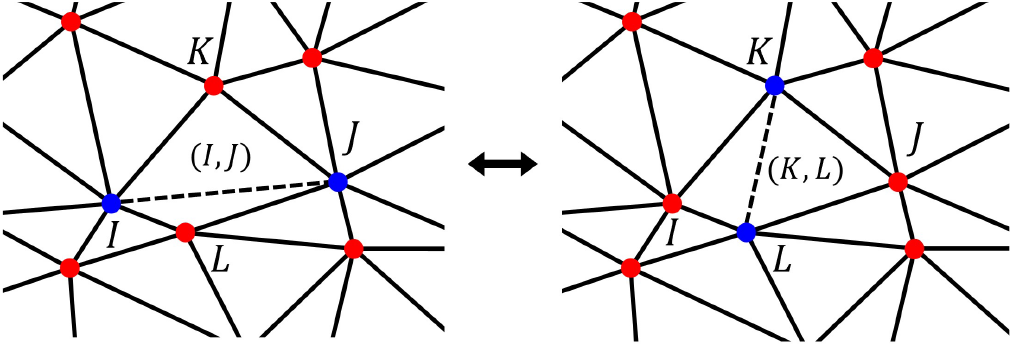
An edge swap in Algorithm 1 in a quadrilateral of cells, *I, J, K*, and *L*. In the forward direction, edge (*I, J*) is deleted while edge (*K, L*) is added.

1. Start the Monte Carlo (MC) simulation with the system in state *γ* defined by the triangulation *𝒯*.
2. Select an edge, (*I, J*) ∈ *𝒯*, in the cell center triangulation, uniformly at random (see Fig. 6).
3. Find the common neighbors, *K* and *L*, for the edge (*I, J*) (see Fig. 6). Note that the algorithm ensures that, at any time, two neighboring cells have exactly two common neighbors.
4. If the edge (*I, J*) yields a ‘valid’ swap (see SI section 3), attempt a T1 transition by removing the edge between centers *I* and *J* and adding an edge between centers *K* and *L*, proposing a new state *γ*′.
5. Calculate the Hamiltonian (or energy gain) *H*_*γγ*_*′* (defined in Eq. 16) in the swapping *γ → γ*′.
6. Accept or reject the transition *γ → γ*′ based on the transition probability 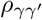, defined as:

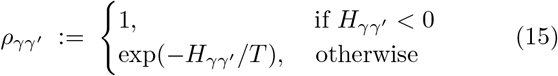

where *T*, the so-called temperature, models the magnitude of fluctuations in the cellular junctions. Note that in every successful swap, one edge is replaced by another, resulting in the total number of edges, say *M*, remaining conserved.
7. Repeat steps 2 − 6 above for *N*_*MC*_ Monte Carlo iterations. A pseudocode for the above algorithm is presented below.

##### Algorithm 1 Updating network topology

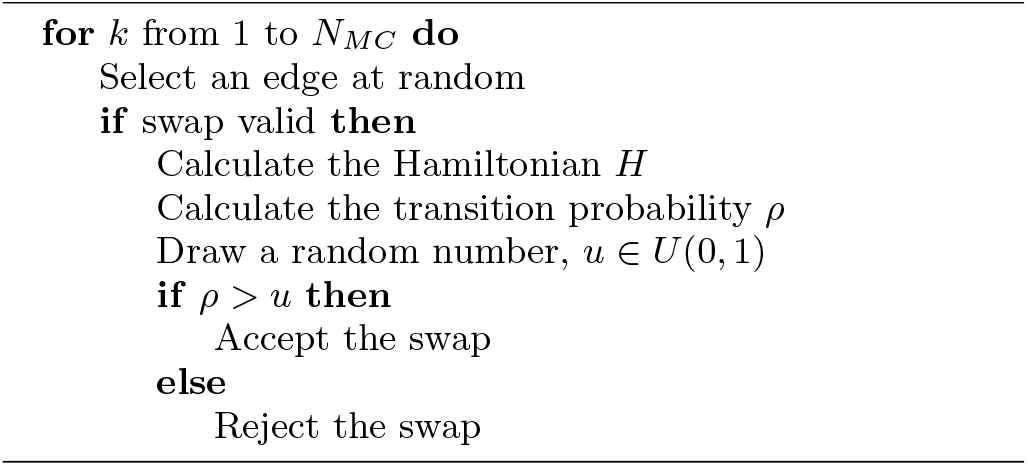

Each MC iteration corresponds to attempting a T1 transition for a set of four cells (*I, J, K, L*). The expression for Hamiltonian 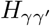 (referenced in Eq.(15)) is motivated thermodynamically by the kinetics of T1 transitions [42]. A T1 transition between states *γ* and *γ*′ involves a transition state, say *γ*_*T*_, defined by a four-fold vertex where cells *I, J, K*, and *L* intersect (see Fig. 7(a)). In the MNM, this vertex is located at the centroid of the convex quadrilateral formed by ***x***_*I*_, ***x***_*J*_, ***x***_*K*_, and ***x***_*L*_. Let 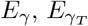, and 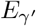 be, respectively, the internal energies of states *γ, γ*_*T*_, and *γ*′. The Hamiltonian 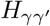 for the transition *γ → γ*′ is now defined as:

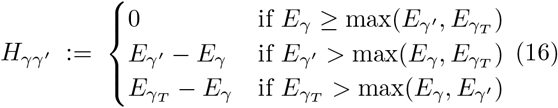

**FIG. 7:**
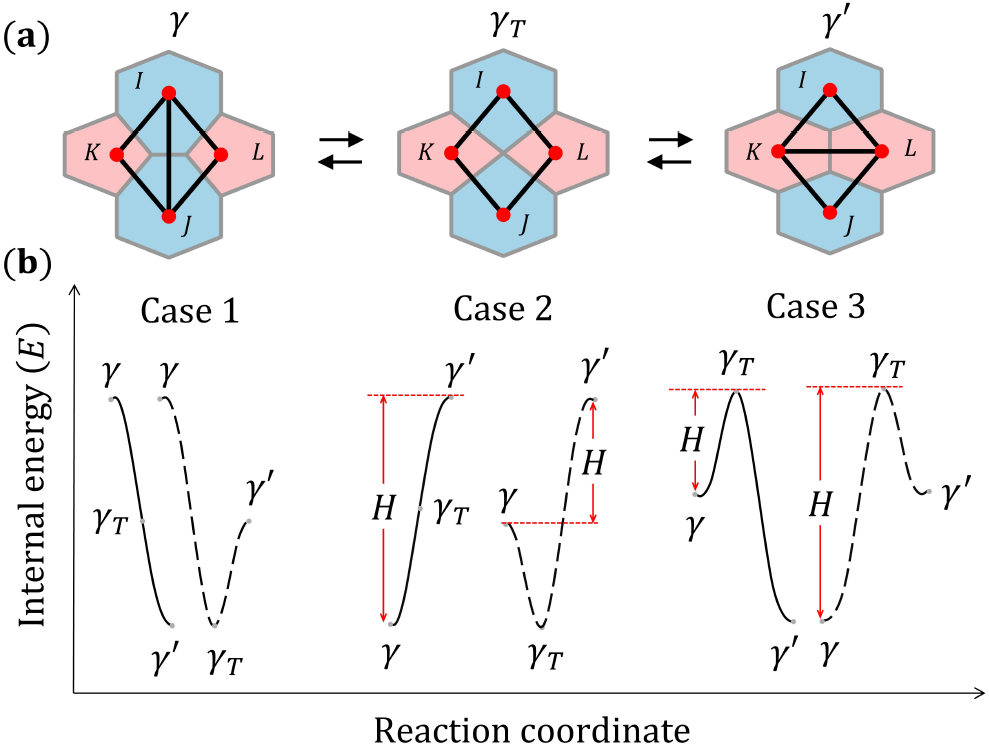
(a) The cellular configuration and associated cell center network during a T1 transition between state *γ* and *γ*′ via the transition state *γ*_*T*_; (b) Potential energetic profiles of a T1 transition for various case scenarios.

The above definition accounts for all possible energy landscapes for the states (*γ, γ*′, *γ*_*T*_) involved in the T1 swapping (see Fig. 7(b)). Typically, the transition state *γ*_*T*_ is unstable (i.e., *E*_*γ*_*T > E*_*γ*_, 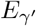), but there could be instances where the four-fold vertex is stable [43]. For simplicity, the MNM currently assumes that the system cannot persist in the transition state (i.e., *γ*_*T*_ does not belong to the space of accessible states). While this feature can be incorporated into the MNM in the future, cellular configurations in this study consist of only threefold vertices. Consequently, the number of edges, *M*, in the network always remains constant. Furthermore, unless otherwise stated, the internal energy *E*_*γ*_ is taken to be the total stored elastic energy *e* (Eq. (4)). T1 swaps with transition probability 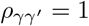 (i.e., 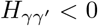) are spontaneous, always lowering the system’s internal energy. Such transitions are effectively barrierless. In contrast, accepted swaps with 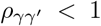 (when 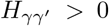) arise from junction fluctuations (*T >* 0) and can be viewed as entropically driven.

By construction, the Markov chain generated by Algorithm 1 is finite (when *N*_*cells*_ is finite), irreducible, and aperiodic. Because it also satisfies the detailed balance condition (shown in Appendix D), it converges to a unique stationary distribution. Let us now study the evolution of the system during a MC simulation step. Note that the MC simulation does not involve a physical time. Hence, this ‘evolution’ refers to the MC iterations. Starting with a random (Poisson) distribution of cell centers and their corresponding Delaunay triangulation, we allow the triangulation to update according to Algorithm 1 for different values of temperature *T*. Convergence to an equilibrium state is shown in Fig. 8, where we plot energy per cell (*e/N*_*cells*_) versus MC iterations. The nature of equilibrium is dependent on *T*. At very low temperatures (*T* ≈ 0), the algorithm becomes trapped in a local minima, accepting only energetically favorable (barrierless) swaps and limiting the exploration of the state space. In this case, the system attains a state of static equilibrium, and rearrangements eventually cease to occur. A slightly higher temperature, *T* = 0.05, enables a broader exploration of the state space, resulting in a lower energy minimum. Further high temperatures, *T* = 0.2 and *T* = 0.5, inject sufficient energy into the system, leading to a dynamic equilibrium at a higher energy level. In the cases *T >* 0, the system fluctuates around the stationary distribution, and fluctuation-induced swaps obscure the underlying stationary distribution. That said, high-temperature regimes can still help model biological phenomena that exhibit significant levels of noise, as we will see later in section III. We reiterate that in this illustration, the term ‘equilibrium’ is used with respect to the number of MC iterations.

**FIG. 8:**
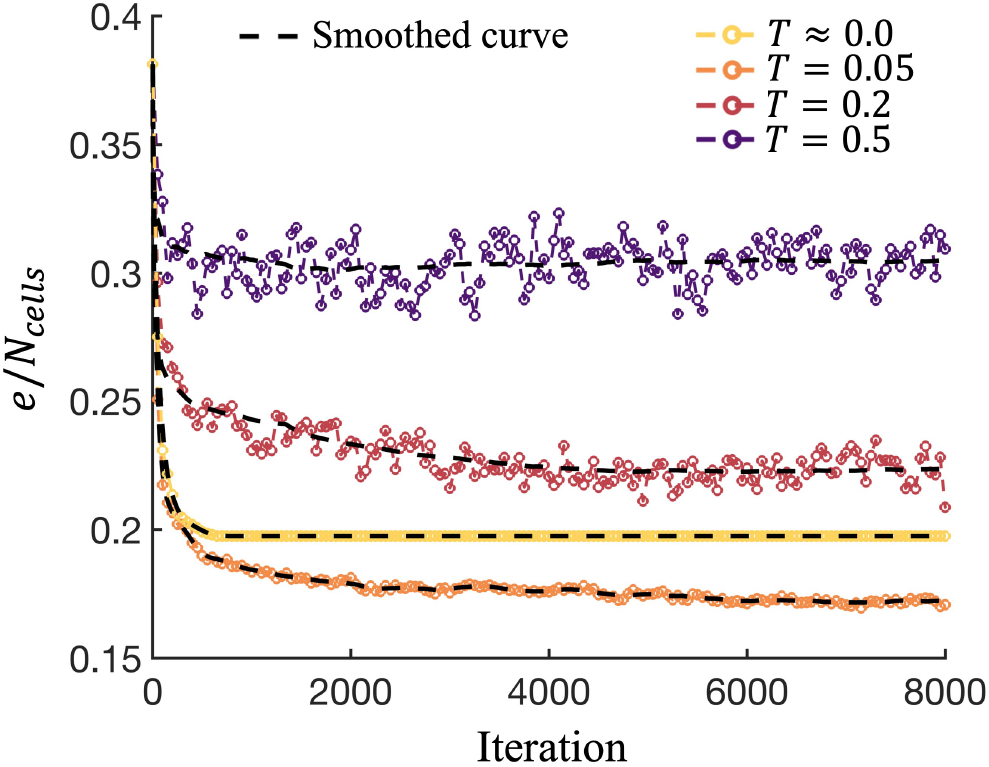
Energy per cell 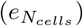 during Algorithm 1 for different temperatures, *T*. The initial configuration (shown later in Fig. 9(a)) is given by a Poisson distribution of cell centers (initialized with the Delaunay triangulation). Simulation parameters: *k*_*p*_ = 0.1, *p*_0_ = 3.

We emphasize that the above-described MCMC implementation of cell rearrangements is highly flexible, in the sense that the Hamiltonian, *H*, can be modified to account for many additional features. For example, here, we do not tie T1 transitions directly to junction length; thus, T1 transitions can appear discontinuous. However, to make cell rearrangements more continuous, one could add a term to the Hamiltonian that penalizes T1 transitions with long junction lengths (similar to VDMs).

### D. The Split-Step Framework

We see from sections II B 1 and II C 1 that while cell center positions are updated based on true dynamics (Newton’s laws of motion), network topology is updated through auxiliary dynamics (MCMC), i.e., no physical time scale is inherent to the Metropolis scheme (Algorithm 1). To introduce a rearrangement timescale, we set a frequency *µ* at which rearrangements are implemented relative to the physical timescale. For instance, *µ* = 100 implies cells are allowed to rearrange only at time steps *t*_*n*_ = 100, 200, 300, … and so on, until the total number *N*_*ts*_ of time steps. The characteristic time, *τ*_*T* 1_, and consequently the rate, *k*_*T* 1_ ~ 1*/τ*_*T* 1_, of T1 transitions in the MNM can then be expressed as:

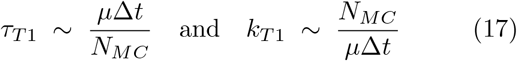

Note that *µ* is dimensionless by definition. The complete implementation of the MNM is now described in Algorithm 2. In practice, we do not perform a sufficiently large number (*N*_*MC*_) of MC iterations for the system to reach equilibrium (see Fig. 8) within every invocation of Algorithm 1 (or MC simulation), as this would effectively yield a quasi-static simulation with respect to T1 transitions. By performing a limited number of MC iterations, we can ensure that the system evolves dynamically over time. To estimate *N*_*MC*_, it would be convenient to express it in terms of the fraction *f* of edges that can undergo swaps within a single MC simulation run. Recalling that the total number *M* of edges in the network remains conserved, we set *N*_*MC*_ = *f M* (rounded off to the nearest whole number, where *f ∈* [0, 1]). The timescale *τ*_*T* 1_ and rate *k*_*T* 1_ can then be rewritten, respectively, as *τ*_*T* 1_ *~ µ*Δ*t/f* and *k*_*T* 1_ *~ f/*(*µ*Δ*t*).

#### Algorithm 2 MNM implementation

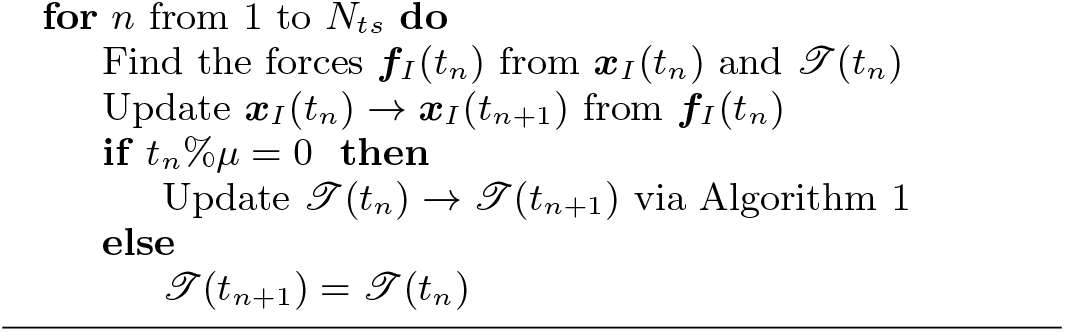

It can be seen that, in the MNM, two distinct mechanisms operate simultaneously with timescales governed by *η* (for center motion) and *µ* (for T1 transitions). The existence of two different timescales in the MNM critically allows us to model the tissue dynamics governed by cell motion and cell rearrangements independently.

*Note*. There could be instances where cell center motion may interfere with the validity of the given cell center triangulation. Thus, we must impose some constraints on cell center motion, addressed in detail in section 3 of the SI.

#### 1. Illustration

To illustrate the methodology outlined in this section, let us now study the dynamics of a monolayer with *N*_*cells*_ = 576. The starting dimensions, *L*_*x*_ and *L*_*y*_, of the rectangular periodic domain are chosen such that *L*_*x*_ *×L*_*y*_ = *N*_*cells*_, and hence ⟨ *a*_*I*_⟩ = 1 (where ⟨·⟩ denotes the mean over all the cells in the population). During the simulation, the boundaries are allowed to relax (i.e., the averaged monolayer stress *σ*_*xx*_ = 0, *σ*_*yy*_ = 0), using the barostat. For simplicity, let us assume no thermal fluctuations (i.e., *T* ≈ 0). To illustrate how the MNM decouples both the mechanisms − cell center motion and cell rearrangements −, we consider two cases: (1) only cell center motion (i.e., *µ→ ∞ ⇒ k*_*T* 1_ = 0, in the MNM, this corresponds to *µ > N*_*ts*_), and (2) only T1 transitions (i.e., *η→ ∞*). For each case, we simulate the monolayer evolution with two different starting configurations: (a) slightly distorted hexagonal lattice (see Fig. 9(a)), and (b) Poisson distribution (see Fig. 9(b)). The cells are color-coded based on their aspect ratio (AR), defined as the minor-to-major axis ratio (see Appendix B). The results are plotted for perimeter elasticity *k*_*p*_ = 0.1 and target shape index *p*_0_ = 3. To quantify the dynamics, we plot the mean shape index, *q* = ⟨ *q*_*I*_ ⟩, and energy per cell, *e/N*_*cells*_, versus time *t* in Fig. 9(c).

**FIG. 9:**
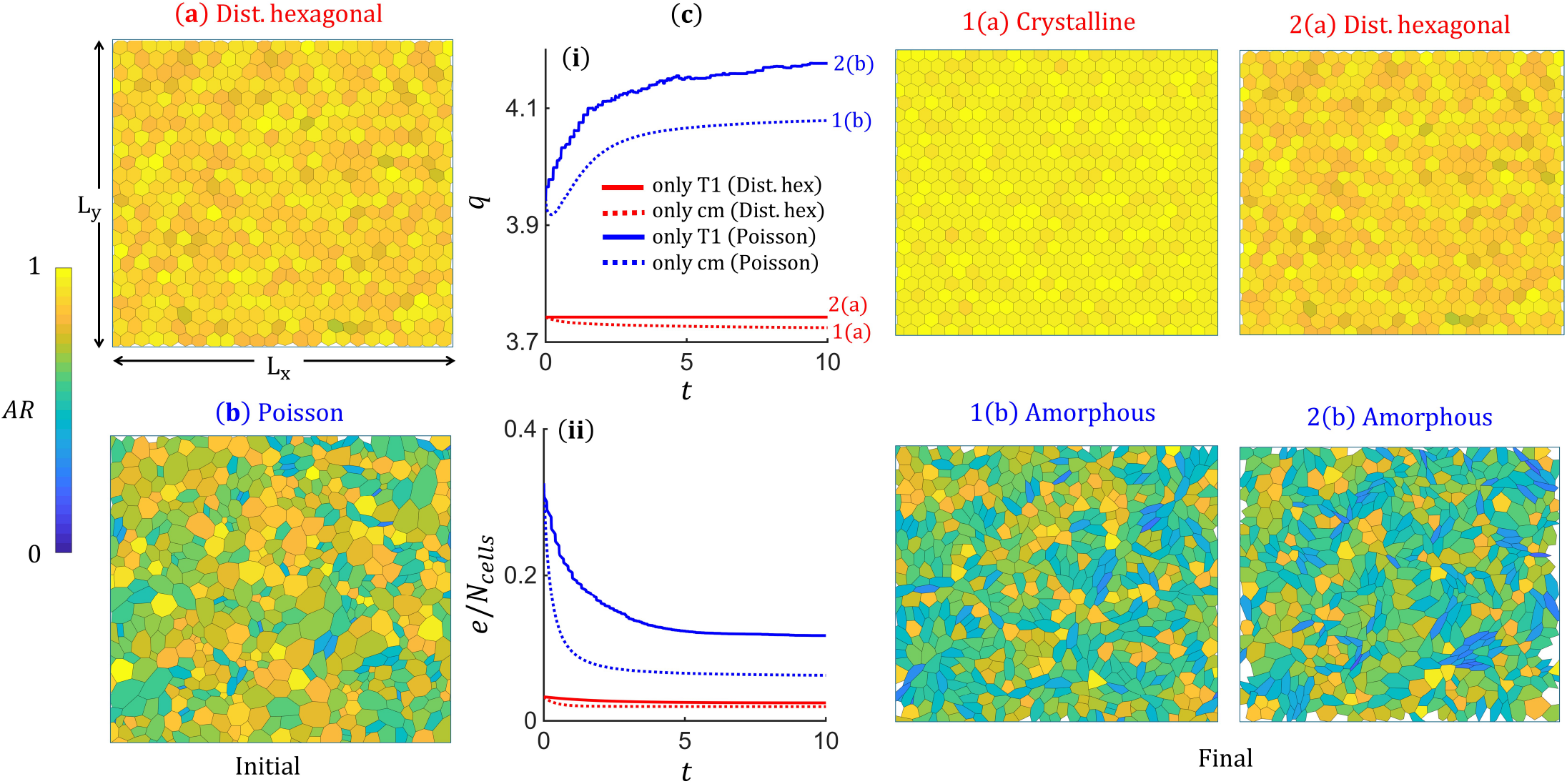
Monolayer evolution within MNM starting from (a) Distorted hexagonal lattice and (b) Poisson (disordered) lattice of cells; Time evolution of (c) (i) Mean shape index *q* and (ii) Energy per cell *e/N*_*cells*_. The final converged monolayer configurations are shown for each case. Simulation parameters: *k*_*p*_ = 0.1, *p*_0_ = 3, *f* = 0.1, *µ* = 10, *η* = 1, Δ*t* = 0.001.

We observe that for all case scenarios, the monolayer evolves to a steady state, where the final configuration (see Fig. 9) depends on both the operating mechanism and the initial configuration. In the case where T1 transitions are absent and the monolayer evolves via the cell center motion, the distorted hexagonal lattice assumes a perfect-crystalline state (1(a)), whereas the random (Poisson) lattice assumes an amorphous state (1(b)). On the other hand, when cell center motion is suppressed and cells can only rearrange, it can be seen that the distorted hexagonal lattice does not evolve (except for boundaries and cells shrinking a little due to barostatting). This is simply because the energy barriers for T1 transitions in this situation are significant, and in the absence of any fluctuations, these energy barriers cannot be overcome. Consequently, the monolayer configuration remains unchanged (2(a)). Interestingly, for the randomly distorted (Poisson) lattice, the monolayer evolves via cell rearrangements, dissipating stored energy through T1 transitions, and converges to a steady amorphous state (2(b)). This is because the cells in the disordered initial configuration are in a high-energy state, resulting in barrierless T1 transitions.

Through this example, we aim to highlight the primary distinction between the MNM and the VTM. Unlike the VTM, the tissue configuration in the MNM can evolve solely through cell-center motion without triggering any neighbor exchanges. At the same time, cell-center motion is not required for T1 transitions as they are governed by an energy-based criterion. This raises the following question. Under what conditions does such behavior arise? We hypothesize that cell-center motion is predominantly governed by substrate friction *η*_*cs*_, whereas cell rearrangements are primarily regulated by tissue viscosity *η*_*cc*_. Because different mechanisms control these two viscosities [27], their magnitudes can differ substantially. Consequently, the characteristic timescales of cell motion and rearrangements would be significantly different as well. In cases where tissue viscosity is high, rearrangements are strongly suppressed. Therefore, the only way cells can lower their stored energy is by moving relative to the substrate while maintaining their neighbors. Alternatively, there could be a situation where substrate friction is significantly higher than tissue viscosity. In this limit, cells can only rearrange while having negligible motion with respect to the substrate. Even though these scenarios would need to be validated experimentally, the MNM framework enables the modeling of these regimes by decoupling cell center motion from cell rearrangements.

Another important takeaway from this illustration is that the monolayer within the MNM framework can settle into multiple stable configurations for the same set of biophysical parameters *k*_*p*_ and *p*_0_ (as demonstrated by Fig. 9). Hence, in contrast to other ABMs [13, 18, 31], the target shape index *p*_0_ within the MNM should not be directly related to the nature of cell packing in stable and stationary tissue configurations. Furthermore, the existence of the tissue in both crystalline and amorphous states for the same value of *p*_0_ should not be interpreted as the unjamming behavior reported in the ABM literature. We now explore this in more detail.

#### 2. Fluctuation-induced unjamming (fluidization)

Fluidization, also referred to as unjamming, in epithelial and endothelial tissues, is one of the most studied phenomena, owing to its importance in collective cell migration and other developmental processes. In this regard, vertex models have been particularly useful, as they directly relate model parameters to tissue mechanical properties. Staple et al. [31] showed that an unjamming transition exists at 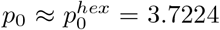, where the shear modulus of the tissue vanishes for *p*_0_ *>* 3.7224. Based on this study, the amorphous (or disordered or floppy) phase of the tissue is deemed fluidized (unjammed), and the crystalline (perfect hexagonal) state is deemed jammed. Later, Bi et al. [18] reported a density-independent unjamming transition, at 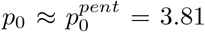, based on the energy barriers of T1 transitions. They found that for *p*_0_ < 3.81, the energy barriers are always finite, and some amount of work must be added to rearrange the cells. Hence, they concluded that tissue exists in a solid-like state for *p*_0_ < 3.81 and in a fluidized state for *p*_0_ *>* 3.81. Unjamming transition at *p*_0_ ≈ 3.81 was also reported in Voronoi-based studies [13]; however, it was later shown that Voronoi models do not exhibit an unjamming transition [44].

Tissue unjamming or fluidization is usually described by a vanishing shear modulus in the disordered/amorphous state. Since the target shape index *p*_0_ in existing ANMs is a direct measure of ordered versus disordered (or floppy) states, *p*_0_ has remained a crucial parameter for quantifying the unjamming behavior. However, there have been some concerns regarding the use of *p*_0_ as a metric for tissue unjamming. Vertex models predict that an increase in cell-cell adhesion, characterized by an increase in *p*_0_, leads to tissue fluidization. According to Bi et. al [18], this observation suggests that adhesion acts differently in dense particulate matter and confluent tissues. Hence, the exact role of intercellular adhesion in tissue fluidization remains a topic of debate and is still under discussion [38].

In the MNM, fluidization is not solely tied to cell-cell adhesion; instead, it is defined dynamically. The most common mode of fluidization in non-proliferating confluent tissues is by T1 transitions. As illustrated in Fig. 9, regardless of whether the converged state is crystalline (ordered) or amorphous (disordered), the tissue is said to be in a jammed state as it attains a steady-state (or static equilibrium). Furthermore, the tissue can exist in the disordered state for 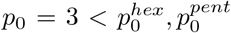 (as can be seen from Fig. 9). Therefore, neither a disordered state nor the shape index parameter *p*_0_ alone is sufficient to conclude that a tissue is fluidized. In fact, from a physical standpoint, the natural way to define the fluidized state of tissue is when it exhibits persistent cell rearrangements (i.e., T1 transitions). In the MNM, a tissue is truly fluidized only when cells continue to rearrange indefinitely. This is achieved through junction fluctuations, with *T >* 0, such that cells are sufficiently excited to regularly cross the energy barrier for rearrangements (Fig. 7). In the last illustration, the system was simulated at *T* ≈ 0, leading to jamming of the cells once the barrierless T1 transitions ceased.

To demonstrate unjamming behavior within the MNM, we now repeat the simulation in section II D 1 and Fig. 9, but this time incorporating both cell motion and cell rearrangements, including those driven by fluctuations (i.e., *T >* 0). The results are plotted for the starting configuration given by the random (Poisson) distribution (see Fig. 9(b)). To quantify fluidization, we plot, in addition to the mean shape index *q* (Fig. 10(a)), energy per cell *e/N*_*cells*_ (Fig. 10(b)), and the number *N*_*T* 1_ (Fig. 10(c)) of T1 transitions versus time *t*. We see from Fig. 10(c) that the monolayer with *p*_0_ = 3 exhibits persistent T1 transitions for *T >* 0, with *N*_*T* 1_ increasing with *T*, thereby indicating unjamming. While the system energy monotonically relaxes, approaching a minimum (a dynamic equilibrium state) for all values of temperature *T* considered (see Fig. 10(b)). The mean shape index, throughout the simulation, is higher for higher temperatures. We find that with fluctuation-induced cell rearrangements and cell motion working in conjunction, we can drive a random (Poisson) initial distribution of cells toward semi-crystalline states (see Fig. 10(d) where we plot the monolayer configurations at the end of simulation for each temperature), with the degree of disorder increasing with *T*.

**FIG. 10:**
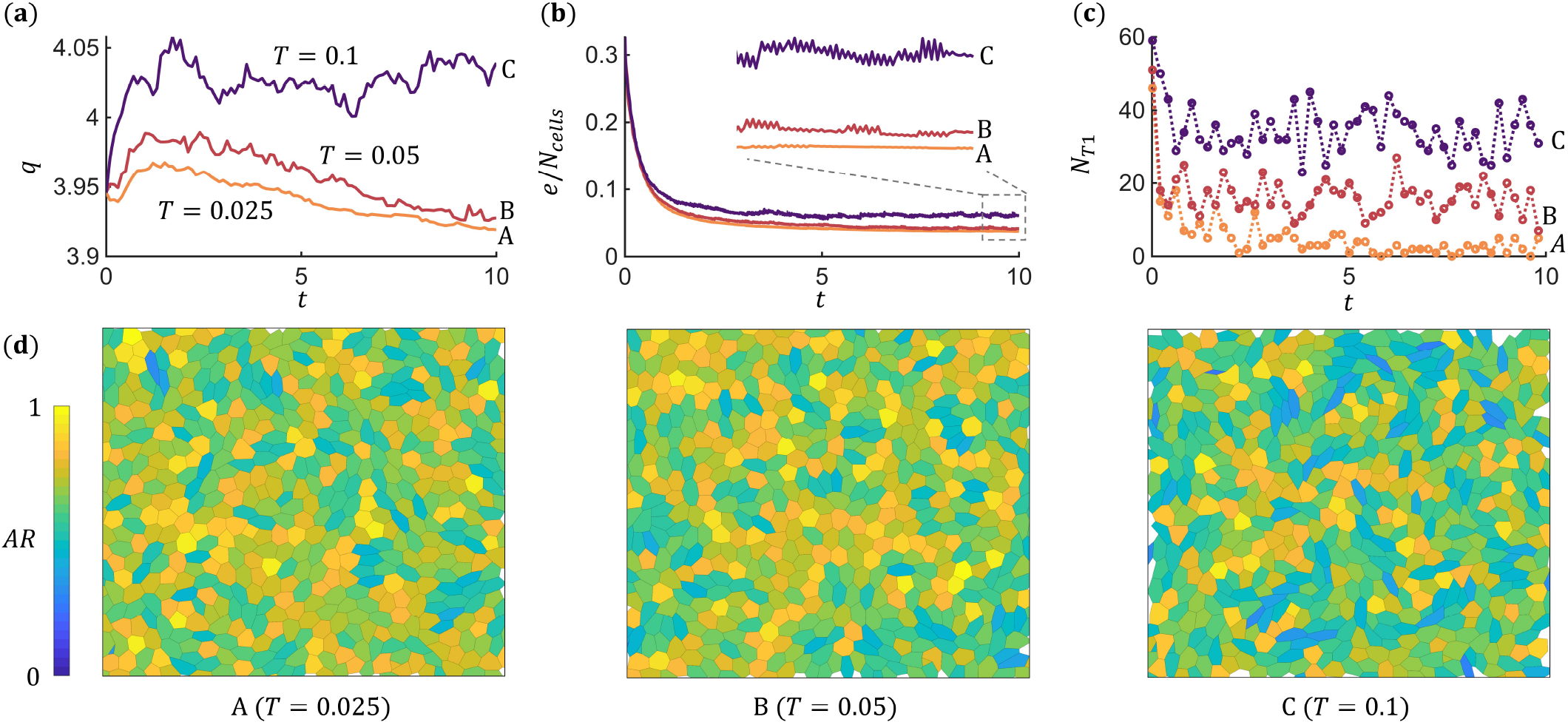
MNM behavior for a Poisson initial condition with cell center motion and cell rearrangements in conjunction for different magnitudes of junction fluctuation *T*. The time evolution of (a) Mean cell shape index *q*; (b) Energy per cell *e/N*_*cells*_; and (c) Number *N*_*T* 1_ of T1 transitions. (d) Final monolayer configuration corresponding to each temperature *T*. Simulation parameters: *k*_*p*_ = 0.1, *p*_0_ = 3, *µ* = 10, *f* = 0.1, *η* = 1, Δ*t* = 0.001.

In a nutshell, in the MNM, though *p*_0_ remains a metric for the averaged aspect ratio of cells (with higher *p*_0_ denoting higher *q*), it should not be used as a director indicator for the nature of cell packing, i.e., crystalline/ordered versus amorphous/disordered state of cells. Neither should it be used as an indicator for the unjamming transition. Instead, in the MNM, the terms fluidization/unjamming are associated with persistent cell rearrangements, rather than with amorphous/floppy/disordered tissue configurations. The critical quantity that governs whether or not the tissue is fluidized is the magnitude of junction fluctuations, *T*. In that sense, though *p*_0_ could facilitate T1 transitions based on the energetics (Fig. 7), some level of noise (*T*) will always be required for the monolayer to maintain a fluidized/unjammed state.

#### 3. Viscoplastic response of suspended monolayer

As mentioned earlier in section II B 2, cell aggregates and suspended monolayers are known to relax stresses via T1 transitions over longer time scales [37, 45–48]. We now explore this long-term viscoplastic behavior using the MNM by taking the example of suspended monolayers.

##### Isochoric biaxial loading

Consider a monolayer initially in the crystalline phase (see Fig. 5(a)), with ground state given by *p* = 3.5251, *a* = 0.8968 (corresponding to *k*_*p*_ = 0.1, *p*_0_ = 3). We have seen previously (from Fig. 5(d) in section II B 2) that elastic loading under uniaxial stretching involves an intrinsic relaxation timescale associated with traction-free boundary conditions in the *y* direction. To avoid complexities in analysis, we impose an entirely displacement-controlled loading for this example problem. Hence, we apply an isochoric biaxial deformation given by deformation gradient of the form ***F*** (*t*) = diag *{λ*(*t*), 1*/λ*(*t*) *}* (as shown in ground state configuration in Fig. 11). For further simplicity, we assume no cell center motion under internal forces, i.e., *η → ∞*. Consequently, the cell centers move affinely under a constant engineering strain rate 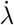 until the deformation reaches *λ* = 2.25, after which the deformation is maintained constant (i.e., 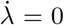). To explore the long-term behavior, we examine two loading rates: (a) fast and (b) slow, relative to the rate of T1 transitions (controlled via *µ*). To characterize the relative loading rate, we employ the dimensionless quantity called the Weissenberg number *W*, defined by the ratio of loading rate 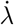 to rearrangement rate *k*_*T* 1_. For this example, we express *W* as *W ≃ µ/N*_*load*_, where *N*_*load*_ is the number of time steps taken from *λ* = 1 to *λ* = 2.25. A value of *W >* 1 thus indicates faster loading compared to the rearrangement rate. Similarly, *W* < 1 denotes the slow loading case. From the mechanics of solids, a suitable scalar measure to characterize the stress state in isochoric biaxial loading is given by the von Mises stress, *σ*_*vm*_, which is calculated as:

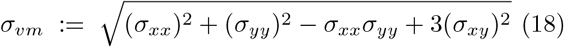

**FIG. 11:**
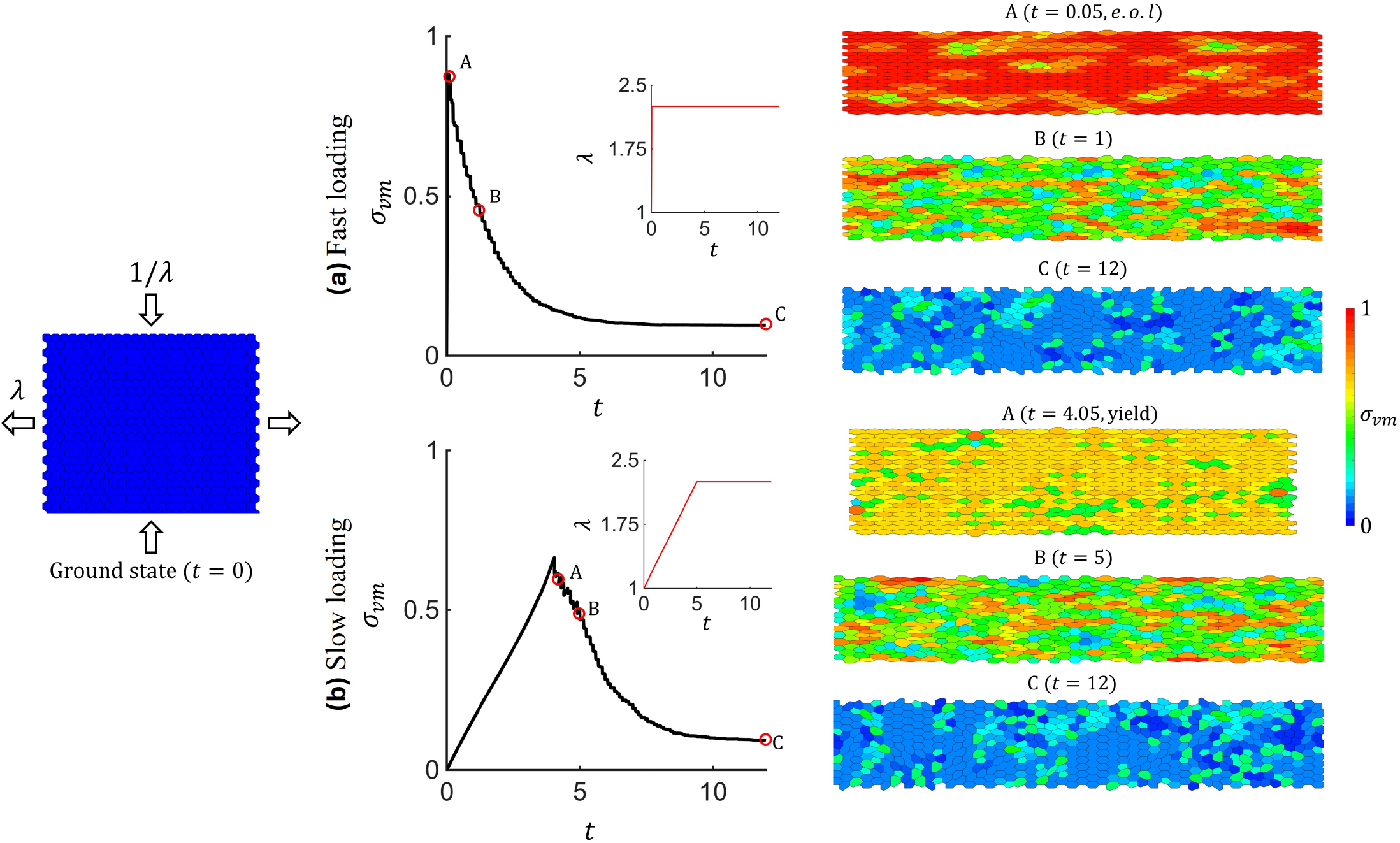
The strain rate-dependent response of a biaxially stretched monolayer with a crystalline ground state, under (a) Fast loading; and (b) Slow loading. The graphs display the time evolution of the global stress state (*σ*_*vm*_), with loading *λ* plotted against time *t* in the inset. The stress-contour plots for monolayer configurations are shown during the relaxation for each case. Simulation parameters: Δ*t* = 0.001, *µ* = 50, *f* = 0.1, *k*_*p*_ = 0.1, *p*_0_ = 3.

Figure 11(a) shows that fast loading (*W >* 1) results in elastic stretching during the loading phase. Once loading is complete (at time *t* = 0.05 or point A), the mono-layer begins to relax through cell rearrangements (see the corresponding monolayer configuration A). In this example, we set the magnitude of fluctuations *T* ≈ 0; hence, all the rearrangements are stress-induced [46]. After the stretch is held constant, the monolayer continues to relax (see monolayer configurations B and C), as cells return to their stress-free hexagonal geometries. Note that cells are color-coded based on their von Mises stress state.

In the slow loading case (*W* < 1), the monolayer begins to relax during the loading phase (i.e., before the loading is completed at *t* = 5 or point B). From Fig. 11(b), it can be seen that the peak stress attained by the mono-layer during loading is reduced in comparison to the fast loading case. That is, the MNM predicts that the mono-layer yields after a certain threshold point (at *t* = 4.05 or point A in this example). The monolayer configuration at the yield point and the end of loading are shown in contour plots A and B, respectively. The final relaxed state (*t* = 12 or point C) is found to be similar for both loading rates.

We notice from the stress contour plots that the mono-layer exhibits spatial heterogeneity in stress distribution. From the contour plot A in Fig. 11(a), we see that relaxation starts at discrete locations, propagating outward. A similar observation is made during yielding (contour plot A in Fig. 11(b)), where yielding occurs in bundles. These observations are reminiscent of force chains, a common emergent phenomenon in granular or particulate media [49]. In granular media under external stress, the load is not shared evenly but is transmitted along pathways where the contact between the particles is higher. These pathways, known as force chains, bear the majority of the applied load. In MNM monolayers, we observe localized regions of low-stress cells where rearrangements have occurred. However, the chains break over the range of a few cells and do not form a continuous network, unlike force chains in granular media. This deviation in behavior can be attributed to the stochastic implementation of cell rearrangements in the MNM. Although the most energetically favorable rearrangement locations are near the yield sites, due to the random selection of potential rearrangement locations (in Algorithm 1), these swaps may not always be selected, resulting in breaking of the chains.

## III. RESULTS

Many biological phenomena involving complex cell geometries require a generalized model, such as the MNM presented here, to accurately capture the morphodynamics. As a model system, we consider the vascular endothelium, the cell monolayer that lines the inner surfaces of blood vessels. The endothelium plays a critical role in maintaining vascular homeostasis. It is central to our understanding of vascular diseases, such as atherosclerosis, and their pathological complications, including heart attacks and strokes. It has been shown that atherosclerotic lesions develop less frequently in arterial regions with elongated and aligned endothelial cells, highlighting the need to understand the shape and alignment behavior of cells in the endothelium [50, 51]. Here we will look at (A) fluidization and emergence of local orientational (nematic) order in the endothelium upon treatment with vascular endothelial growth factor (VEGF), and (B) substrate topography-induced endothelial cell alignment on microgroove substrates [52].

### A. Emergence of nematic order

Experiments show that treating endothelial cell monolayers with VEGF leads to an increase in cell elongation [53]. This can be seen in Fig. 12a(i), with the control (no VEGF) monolayer having a mean shape index *q* = 4.15 and the VEGF-treated monolayer having *q* = 5.12. An increase in average cell elongation (from control to VEGF) directly suggests an increase in target shape index *p*_0_. Physically, an increase in target shape index can be achieved by (1) an increase in cell-cell adhesion *k*_*t*_ or (2) a decrease in contractility *k*_*p*_ or (3) both (see Eq. (5)). The actual mechanism through which VEGF influences monolayer properties is not entirely understood.

**FIG. 12:**
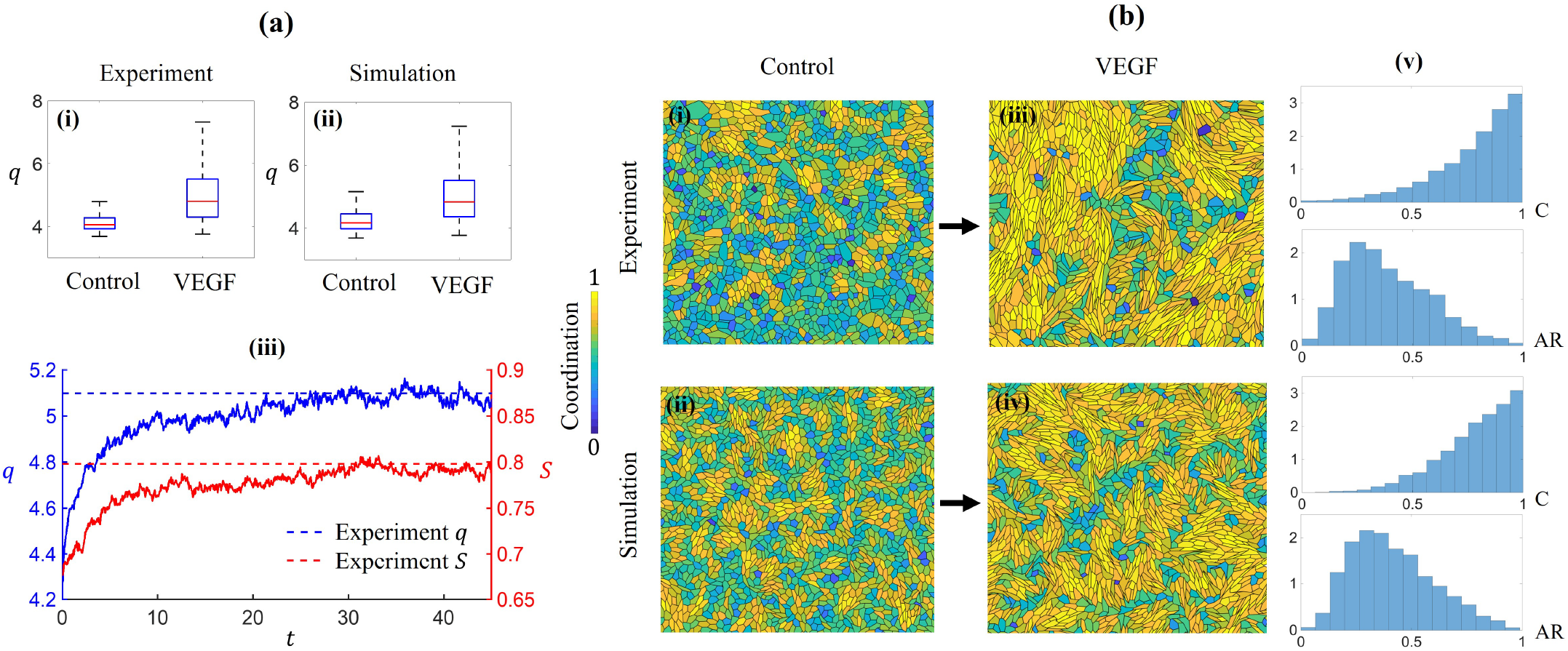
(a) Box plots (outliers omitted) of shape index for the (i) in vitro experiment and (ii) the MNM simulation, showing the control and VEGF states, (iii) morphodynamics of the monolayer within MNM. (b) Processed experimental images for (i) control and (iii) VEGF-treated monolayer and the corresponding snapshots of simulated monolayers in (ii) initial and (iv) toward the end of simulation (*t* = 45) configurations, (v) the histograms with coordination *C* and aspect ratio *AR* for the VEGF state. Histogram data are combined across 3 identical experiments/simulations with *~* 6000 cells total. Simulation parameters: *p*_0_ = 5.12, *k*_*p*_ = 0.1, *T* = 0.5, *µ* = 10, *η* = 1, *f* = 0.1, *f*_*a*_ = 0, *N*_*cells*_ = 2000. For details about experiments, refer to section 4.1 in the SI.

Along with cell elongation, experimental results show an increase in local orientational order, where a cell’s major axis (or director ***u***_*I*_ defined in Appendix B) tends to align with that of its neighbors. Interestingly, the order breaks down over a range of a few cells, resulting in a spatially heterogeneous nematic phase. To quantify this alignment behavior, we define a measure of cell-neighbor coordination, *C*_*I*_ for cell *I*, as:

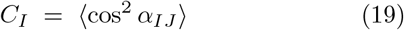

where cos *α*_*IJ*_ = ***u***_*I*_ *·* ***u***_*J*_ and the mean is taken over all the immediate neighbors (those with shared vertices) of cell *I*. A coordination value of *C*_*I*_ = 1 indicates that the cell director is completely aligned with those of its neighbors, whereas *C*_*I*_ = 0 implies no alignment. A global order parameter can also be defined as:

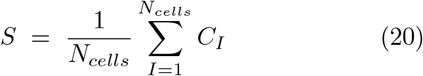

to measure the average coordination of cells in the monolayer. Treatment with VEGF is found to promote the local alignment of cell directors and increase the global order parameter *S* from 0.616 in the control monolayer to 0.795 in the VEGF-treated monolayer (Note that here, we calculated *S* from the post-processed image of the experimental monolayer by fitting cells to polygons). We now aim to understand the emergence of local nematic order in the endothelium, which is essential as it plays a key role in various morphogenetic processes [5, 7, 54].

To simulate the effect of VEGF on the cell monolayer, we first obtain an initial configuration (i.e., the simulation control) that mimics the post-processed experimental control configuration (see Fig. 12b(i), where cells are color coded based on their coordination). The simulation control configuration (as shown in Fig. 12b(ii)) is set such that the shape index distribution (Fig. 12a(ii)) matches that of the experimental control (Fig. 12a(i)). To model the effect of VEGF, the target shape index, *p*_0_, of the monolayer in MNM is set at 5.12 to match the mean shape index of the VEGF-treated monolayer in experiments (Fig. 12a(i)). Other simulation parameters are given by *k*_*p*_ = 0.1 and *T* = 0.5, where the temperature, *T*, is chosen such that the simulation *q* converges to *p*_0_. We note that this requires a very high temperature (*T* = 0.5), indicating high levels of rearrangement activity and, therefore, tissue fluidity. Furthermore, in the simulation, we allow both mechanisms, i.e., cell center motion and T1 transitions, to operate simultaneously (with parameters *η* = 1.0, *µ* = 10 and *f* = 0.1). The monolayer then evolves from *q* = 4.15 to *q* ≈ *p*_0_ = 5.12 and attains a dynamic equilibrium (see Fig. 12a(iii)).

Akin to the experiments, the MNM predicts the emergence of locally aligned clusters of elongated cells. This can be seen from Figs. 12b(iii) (VEGF-treated monolayer in experiments) and 12b(iv) (monolayer configuration at the end of simulation, *t* = 45), in which the coordination behavior in the simulation results matches that of the experiments. Interestingly, when the target shape index *p*_0_ is set to match the experimentally observed mean value of 5.12, the global nematic order parameter *S* also converges towards the experimental value of 0.795 (see Fig. 12a(iii)), further validating the simulation results. For quantitative comparison, we plot the distribution of cell aspect ratio (AR) and coordination (C) from the final simulation snapshot (*t* = 45) along with the experimental VEGF-treated state. The histograms in Fig. 12b(v) show that the MNM results agree closely with the experimental monolayer configuration.

In these results, while the role of cell-center motion cannot be neglected, we observe (not shown here) similar behavior in the case where the motion of cell centers is restricted (i.e., *η → ∞*). This demonstrates that cell rearrangements (via T1 transitions) are the principal driving mechanisms by which fluidized monolayers exhibit nematic ordering. We infer from the model that in a fluidized state, cells tend to rearrange to align with their direct neighbors. This collective behavior emerges naturally within the MNM and is entirely an outcome of thermodynamically motivated T1 transitions (Algorithm 1). This is in contrast to previous studies where nematic ordering was obtained phenomenologically by making junction tension a function of cell shape [19].

Lastly, we note that MNM simulated monolayers tend to be more spatially homogeneous (Figs. 12b(ii, iv)) than observed experimentally (Figs. 12b(i, iii)). This discrepancy can be explained as follows. In the simulations, there is a single target geometry that all cells seek to match; that is, all cells have the same target area and the same target perimeter. However, cell-to-cell differences in VEGF receptor expression have been reported in various studies, suggesting there may be an inhomogeneity in the target shape index of cells in real monolayers. Further efforts could be made to more closely match experimental data by fine-tuning and introducing cell-to-cell variation in model parameters. We leave this aspect for future work.

### B. Topography-induced cell alignment

In the last example, we observed an emergent collective response where cells aligned locally with their neighbors. Here, we model topography-induced alignment behavior of endothelial cell monolayers as observed by Leclech et al. [52]. The objective of the experimental study was to understand how biophysical cues from the underlying extracellular matrix (ECM) influence collective behavior within the monolayer. To mimic the in vivo environment and study the effects of ECM anisotropy, Leclech et al. cultured endothelial cell monolayers on either flat or microgroove substrates that model the anisotropic topography of the vascular basement membrane. They observed that endothelial cells on microgroove substrates aligned and elongated along the direction of the grooves (*x*-axis). This behavior became more pronounced with the depth *d* of the microgrooves, as shown in Fig. 13(a), where the cells are color-coded based on their alignment *θ* (i.e., the angle between cell director ***u***_*I*_ and the groove direction ***n***_*g*_). The behavior on flat substrates (*d* = 0) was isotropic, with cells orienting their director in random directions.

**FIG. 13:**
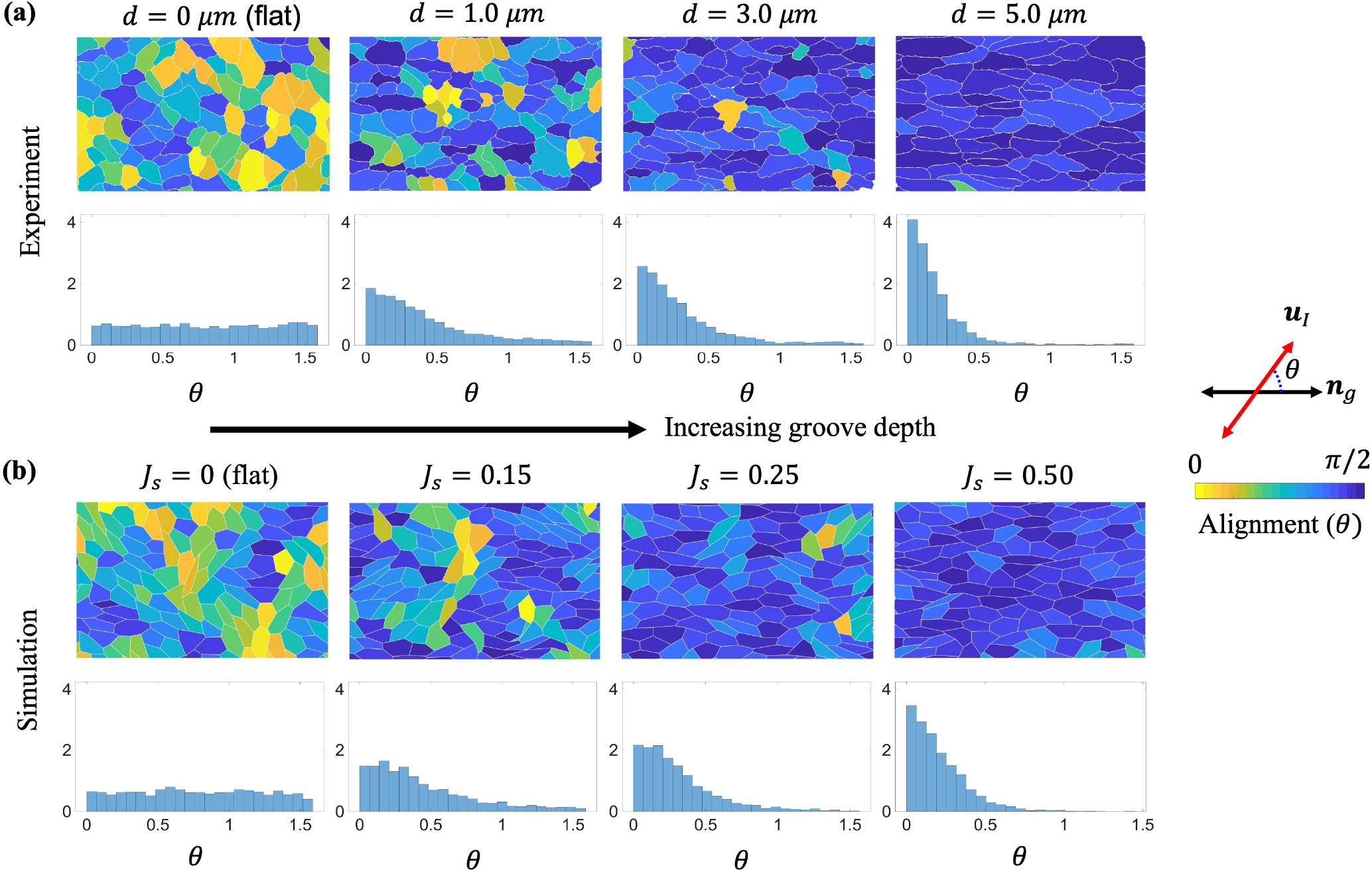
Monolayer renderings and histograms of cell alignment with the groove direction (*x* − axis), *θ*, for (a) increasing groove depth, *d*, (in the experiments) and (b) increasing substrate alignment penalty strength, *J*_*s*_, (in the simulations). Histogram data combined from 3 identical experiments/simulations. Experiments range from 1666-3107 cells per treatment group, while simulations have 2250 cells per treatment group, with *N*_*cells*_ = 750. Simulation parameters: *p*_0_ = 4.4066, *k*_*p*_ = 0.1, *T* = 0.12, *µ* = 10, *η* = 1, *f* = 0.1. For experimental details, refer to section 4.2 in SI.

We now simulate this alignment behavior using the MNM. For the current study, we only focus on the effect of substrate anisotropy on cell alignment, while neglecting active migration (i.e., *f*_*a*_ = 0). The non-trivial effect of grooves on active collective migration will be addressed in a future study. Experimental observations suggest that cells respond to grooved substrates by aligning their directors along the groove direction. To phenomenologically capture this observation, we add a cell-substrate interaction energy contribution, say *e*_*cs*_, to the internal energy, *E*, and write:

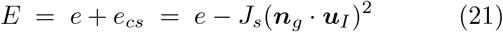

where the penalty parameter *J*_*s*_ (subscript *s* denotes substrate) characterizes the strength of the cell-substrate interaction. The additional energy term penalizes any deviation in the orientation of the cell director from the groove direction, ensuring that cells preferentially rearrange to align their director along the groove direction. We first obtain an initial configuration by running the MNM with a target shape index matching the experimental control (in this case, the flat substrate where *J*_*s*_ = 0 with *p*_0_ = 4.1066), and find a temperature (*T* = 0.12) that achieves convergence of the simulation mean shape index *q* to *p*_0_ = 4.1066. We then increase the alignment strength parameter *J*_*s*_ to model the effect of grooves.

The snapshots for the simulated monolayer toward the end of the simulation are plotted in Fig. 13(b) (for different values of alignment strength, *J*_*s*_), against the experimental images in Fig. 13(a). For quantitative comparison between simulations and experiments, we also plot the corresponding histograms for the alignment angle *θ*. We see that with higher *J*_*s*_, cells increasingly align along the groove direction as expected. By modulating *J*_*s*_, it is possible to match the level of alignment observed in experimental images, indicating that the biophysical parameter *J*_*s*_ represents groove depth. It is essential to mention that the alignment energy term is introduced only in the internal energy *E* (and, as a result, in the Hamiltonian *H*) governing neighbor exchanges. The motion of cell centers remains governed solely by the elastic energy *e* associated with cell shape (see Eq. (4)). While in principle one could additionally include the alignment energy in the elastic energy function, such that the force on a cell center becomes 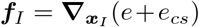, the emergent behavior (akin to the previous example in sec. III A) is dominated by T1 transitions, with a marginal role for cell center motion. The latter, however, is expected to become more important during active migration and is therefore left for future studies.

## IV. CONCLUSION

In this work, we have developed an agent-based model, termed the morphodynamic network model (MNM), to simulate the complex mechanics and dynamics of multicellular systems. In this framework, a cell monolayer is represented as a dynamic triangulation of cell centers (nodes) and neighbor connections (edges). The MNM follows a split-step procedure that combines aspects of dissipative particle dynamics (to model cell center motion) and a discrete-time Markov chain Monte Carlo process, governed by a Hamiltonian *H* and temperature *T* (to model cell rearrangements). Unlike existing models (VDMs and VTMs), our hybrid approach decouples cell motion from topological transitions, enabling better control of tissue mechanics and geometry. For instance, the MNM allows for modeling cells that are immobile yet topologically active, a scenario that is difficult to implement in VDMs and VTMs due to their inherent coupling between motion and T1 transitions. The MNM critically introduces two distinct time scales: one governing cell motion and another governing topological rearrangements. This separation enables more realistic modeling of systems, such as suspended monolayers. Unlike previous cell center-based models [55] that focus primarily on elastic behavior, the MNM extends to highly fluidized regimes with vigorous cell rearrangements and heterogeneous morphologies (appropriate for phenomena such as VEGF-treated monolayers and cells cultured on microgroove substrates). One of the key features of the MNM is the stochastic implementation of T1 transitions. In addition to being thermodynamically motivated, the use of temperature *T* to govern neighbor exchanges naturally incorporates junctional tension fluctuations, contrasting with time-dependent junction tensions used in other models [20, 23]. We demonstrate the ability of the MNM to model emergent collective behavior, such as nematic ordering, by simulating some in vitro experiments on endothelial cell monolayers. The predicted results align well with the experiments, showcasing the practicality of the MNM methodology.

We primarily focused on systems dominated by cell rearrangements, forgoing those with large-scale cell motion. In fact, the constraints on cell center motion (outlined in the SI section 3) prevent cell motion on scales larger than a single cell. In future work, we will relax this constraint and explore systems with active cell motion, leading to large-scale collective migration. We can thus revisit the problem of endothelial cell alignment on microgroove substrates, additionally modeling the formation of antiparallel streams, as reported by [52]. Additionally, we intend to investigate other collective phenomena, such as the formation and dynamics of topological defects, which are known to play a crucial role in morphogenetic processes like cell extrusion [7, 56]. Many real-life morphogenetic phenomena occur in three-dimensional curved shell-like cell monolayers, such as intestinal organoids [57]. Hence, an interesting application of the MNM would be to investigate emergent collective behavior in these geometries, which would require extension of the MNM to three dimensions. This would also enable the exploration of more complex problems in tissue Engineering, where multicellular aggregates are encapsulated within hydrogel matrices [58]. In these systems, the morphomechanics of the cell aggregate is dictated by the mechanical constraint provided by the hydrogel. Such coupled systems have received limited attention in agent-based models, primarily due to the challenges associated with representing the complex mechanical response of hydrogels [59, 60]. The goal of the MNM, in this regard, is to expand the capabilities of agent-based modeling by enabling simulations of complex tissue-level scenarios that involve dynamic interactions between multicellular systems and their microenvironment.

## SUPPORTING INFORMATION

Supplementary Video 1: Stress relaxation for fast loading (associated with Fig. 11(a)).

Supplementary Video 2: Stress relaxation for slow loading (associated with Fig. 11(b)).

Supplementary Video 3: Morphodynamics of VEGF-treated monolayer simulated using the MNM (associated with Fig. 12).

Supplementary Video 4: Morphodynamics of monolayer over microgroove substrate, as simulated using the MNM (associated with Fig. 13 for *J*_*s*_ = 0.5).

## CODE AVAILABILITY

The code for the morphodynamic network model (MNM) is developed within LAMMPS (a molecular Dynamics simulator) [61] and is available upon request from the authors: Prakhar Bandil^1^ and Franck Vernerey ^2^.

## ACKNOWLEDGMENTS

FJV gratefully acknowledges the support of the National Science Foundation, USA, under Award No. 2135057. The content is solely the responsibility of the authors and does not necessarily represent the official views of the National Science Foundation.

## Appendix A: Non-Dimensionalization

The unit of length is given by 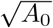, where *A*_0_ is the target cell area (assumed to be the same for all cells). Hence, the area *a*_*I*_ in Eqn. (4) is *a*_*I*_ = *A*_*I*_*/A*_0_, where *A*_*I*_ is the actual area of cell *I*. Similarly, 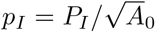, with *P*_*I*_ being the actual perimeter. The position vectors ***x***_*I*_ and ***r***_*v*_ (introduced in Fig. 3) are also normalized and measured in units of 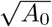. The area *a*_*I*_ and perimeter *p*_*I*_ are calculated as:

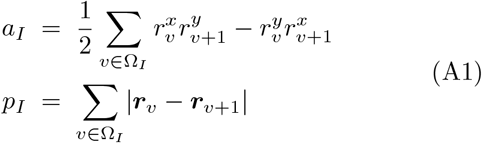

where 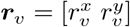. 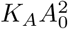 gives the unit of energy *e*, and the unit of force becomes 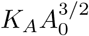, where *K*_*A*_ is the area elasticity (often interpreted as the bulk modulus of the cytoplasm). The damping coefficient *η* is measured in units of tissue viscosity *η*_*cc*_. The unit of time *t*, thus becomes 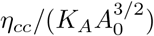. The (normalized) perimeter elasticity *k*_*p*_ and junction tension *k*_*t*_ are given, respectively, by *k*_*p*_ = *K*_*P*_ */K*_*A*_*A*_0_ and 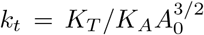. Here, *K*_*P*_ and *K*_*T*_ represent perimeter elasticity and junction tension, respectively, in physical units.

## Appendix B: Cell shape tensor

The major eigen direction for a cell *I* is calculated as follows. We first find the shape tensor ***G***_*I*_ for cell *I* defined as [14, 62]:

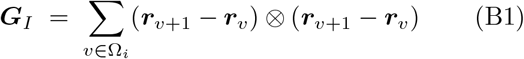

where *⊗* denotes the dyadic product. In matrix form, we can rewrite ***G***_*I*_ as:

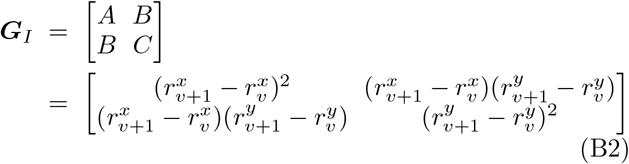

Eigen values, *λ*_1_, *λ*_2_ of ***G***_*I*_, can then be found as:

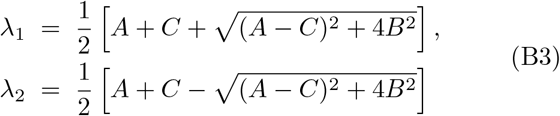

The aspect ratio (AR) for the cell is then calculated as:

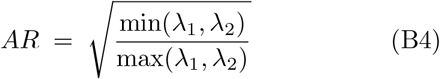

Corresponding to the maximum eigenvalue *λ*_*max*_ = max(*λ*_1_, *λ*_2_), the eigenvector 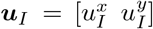, can be determined as:

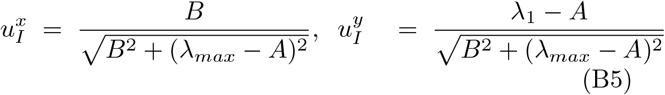

## Appendix C: Ground state

For the crystalline phase where stress-state for each cell is identical (and *a*_*I*_ = *a, p*_*I*_ = *p, l*_*IJ*_ = *l*), we can write (using Eqs. (9), (10), and (11)):

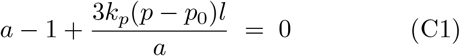

Rewriting the equation only in terms of perimeter (substituting 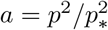 and *l* = *p/*6), yields a cubic function in perimeter, agreeing with [36]:

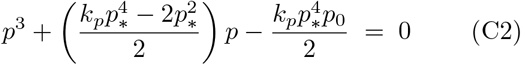

## Appendix D: Detailed Balance

Detailed balance is defined by the condition [41],

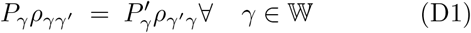

where 𝕎 is the space of all possible states (for a given distribution 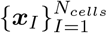 of cell centers). At equilibrium, the probability *P*_*γ*_ of the system existing in state *γ* is given by [41]:

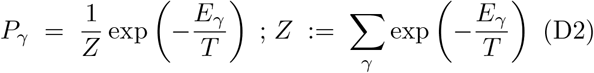

*Z* is the partition function. Checking the detailed balance, we find, for each case in Eq. (16):

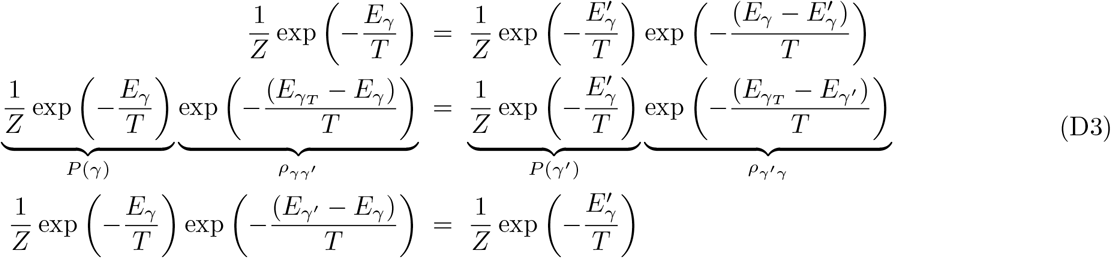

## SUPPLEMENTARY INFORMATION

### 1 Barostatting

For in silico experiments (whether it is stretching of a solid-like monolayer or studying a fluidized monolayer), we need to ensure the monolayer is unconstrained, which is given by ***σ*** = **0**. Note that ***σ*** is the mean stress (or global stress) in the monolayer. A tissue configuration obtained using random (Poisson) distribution of points in a 2D space is generally out of equilibrium as it does not satisfy ***σ*** = **0**. Therefore, the first task is to bring this monolayer to a stress-free state, which will then serve as the starting configuration for our computational experiments. Since the model is coded in LAMMPS, we make use of the barostat feature that deforms the system to attain a desired pressure (In LAMMPS, pressure is used to denote stress tensor). We implemented a simple barostat given by (https://docs.lammps.org/fix_deform_pressure.html):

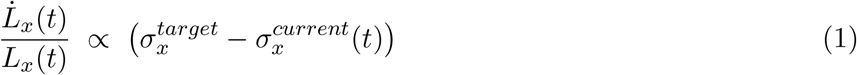

An analogous equation can be written corresponding to the *y* dimension. The dimension of the simulation box is then deformed as:

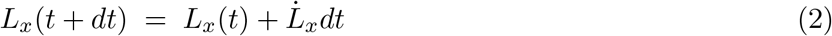

The dilation or stretch in that direction is then:

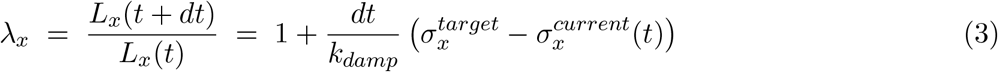

The damping factor *k*_*damp*_ will dictate how fast the system will attain its target pressure. Too small values can result in large fluctuations in the dimensions, whereas too large values would take a long time for the system to attain the target value. Choosing an appropriate value depends on the specific simulation and system. In our study, for Δ*t* = 0.001, we chose *k*_*damp*_ between 1 and 20. This resulted in smooth relaxation.

### 2 Non-affine motion

When the initial configuration is crystalline (symmetric), the motion of cell centers is strictly affine. We now investigate the role of non-affine motion in the elastic response of tissues. We deform an amorphous tissue, resulting in non-zero forces on cell centers and non-affine motion. To begin, we first obtain the ground state of the amorphous tissue. We start with a pseudo-initial configuration given by the Poisson distribution of points with the Delaunay triangulation. In this configuration, the periodic domain boundaries (*L*_*x*_, *L*_*y*_) were set such that ⟨*a*_*I*_⟩ = 1. The stress state in this configuration is found to be *σ*_*xx*_ = *σ*_*yy*_ = 0.66 (corresponding to *k*_*p*_ = 0.1, *p*_0_ = 3). To obtain a stress-free configuration, we allow the boundaries to relax (via barostatting) such that the stress components, *σ*_*xx*_ and *σ*_*yy*_, reduce to 0 (see Figs. 1(a)(i) and (a)(ii) for *t* < 0, which denotes initial relaxation). We note that *σ*_*xy*_ ≈ 0 throughout. The ground state for an amorphous tissue also requires ***f***_*I*_ = 0 for all *I*, which we achieve by running the simulation under passive forces (*f*_*a*_ = 0) to a stationary state. We plot, in Fig. 1(a)(i), the magnitude of averaged force *f* = ⟨|***f***_*I*_|⟩ and observe that it relaxes to zero, indicating a fully relaxed (ground) state (at *t* = 0). This is the true initial configuration as shown in Fig. 1(b)(i).

**FIG. 1:**
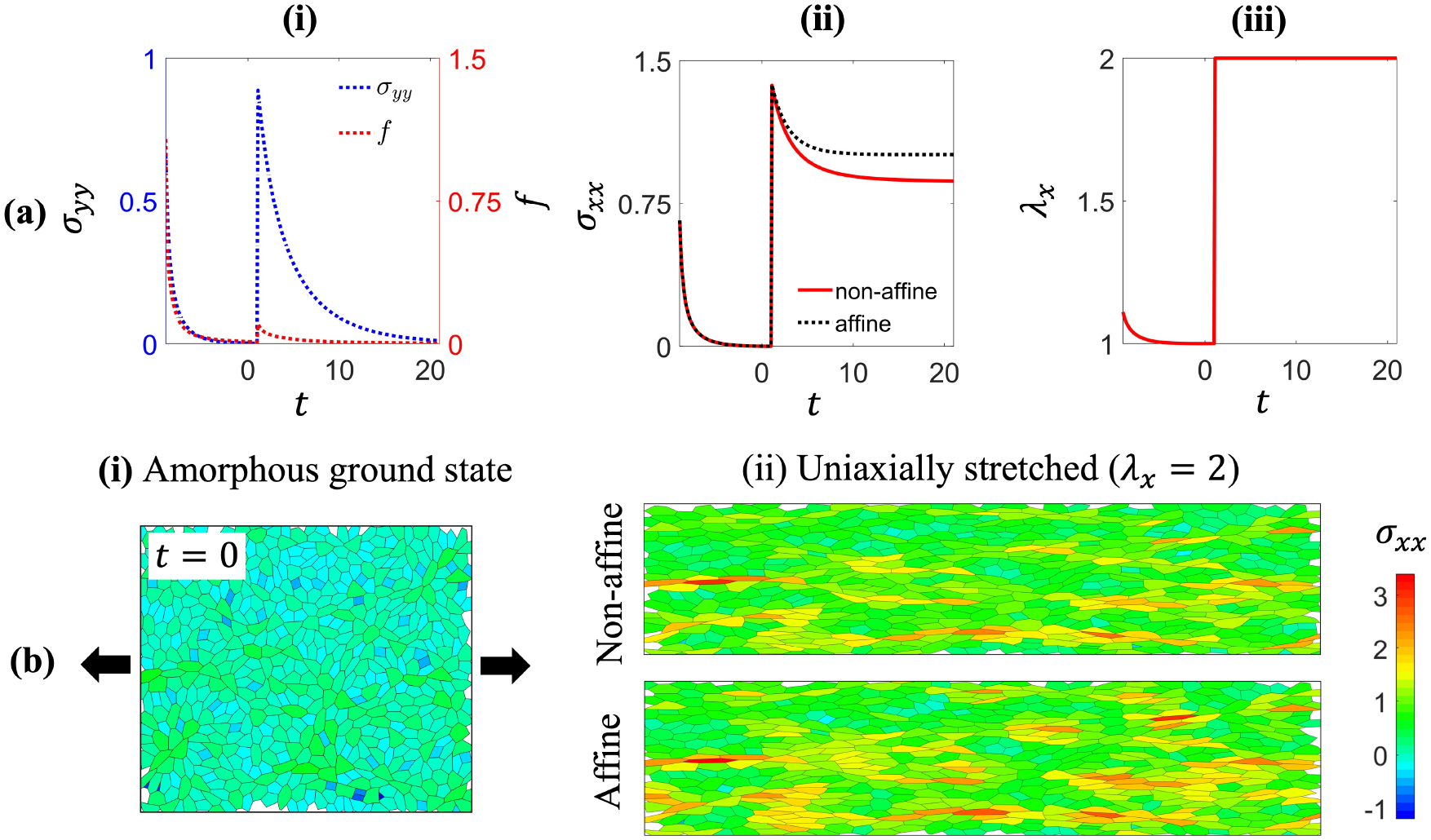
(a) The elastic response of a monolayer under uniaxial stretch with non-affine motion; (i) relaxation of stress *σ*_*yy*_ and average force *f*; (ii) evolution in uniaxial stress *σ*_*xx*_ for affine versus non-affine; (iii) the loading, applied in a single step starting at *t* = 0. (b)(i) Relaxed-initial configuration of the amorphous monolayer; (ii) the stretched configurations for both the affine and non-affine cases.

Note, individual cells are not stress-free. The amorphous ground state exhibits a complex cellular stress distribution, often referred to as residually stressed. The global stress-free state, ***σ*** = **0** (or mechanical equilibrium) is achieved by some cells being under compression while others are in tension. Stretching (with *λ*_*x*_ = 2 as shown in Fig. 1a(iii)) the residually-stressed monolayer uniaxially, we record the elastic response for the affine and non-affine cases. We observe from the stress response plot of *σ*_*xx*_ versus time (Fig. 1a(ii)) that the non-affine case results in a lower equilibrium stress for the same applied stretch. This is because the cells dissipate energy (consequently, stresses) when allowed to move under passive forces. Stress dissipation is also evident from the contour plot in Fig. 1b(ii). Thus, non-affine motion results in a softer elastic response, as is commonly observed Staddon et al., 2023.

### 3 Network regularization

As our model imposes few fundamental restraints on cell shape, as opposed to highly constrained Voronoi models, it is necessary to implement some computational constraints on T1 transitions and cell center movement to maintain biologically valid cell geometries as the monolayer evolves in time. In generalized cell center triangulations, T1 transitions can result in complex configurations that are not biologically feasible, such as overlapping or self-intersecting cells. To prevent nonphysical configurations while maximizing model freedom, we forbid problematic configurations as they arise in the T1 transition algorithm.

1. We require each cell center to maintain at least four neighbors (resulting in cells that have four or more sides). Triangular cells are generally poor approximations of real cells, and in forbidding them, we prevent the potential generation of two-sided cells, which are entirely non-physical.
2. We also require a quadrilateral of cell centers to be convex for a T1 transition to be considered. A T1 transition on a non-convex quadrilateral of cell centers results in an invalid triangulation.

If a T1 transition is proposed that triggers one of the above cases, the swap is rejected. In terms of the rearrangement Hamiltonian, these forbidden states are considered to have infinite energy. Thus, the transitions never occur according to Algorithm 1.

Another class of constraints is on the motion of cell centers or nodes.

1. To prevent cell centers or nodes from overlapping, we apply soft repulsive forces between cell centers. We use the same soft repulsive force as outlined in Barton et al., 2017,

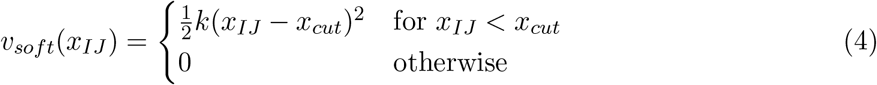

where *x*_*IJ*_ = |***x***_*I*_ − ***x***_*J*_ |. We select *x*_*cut*_ = *l*_0_*/*2.5, where 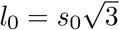 is the distance between cell centers in the regular hexagonal lattice corresponding to a given side length *s*_0_. The resulting force on cell *I* is thus,

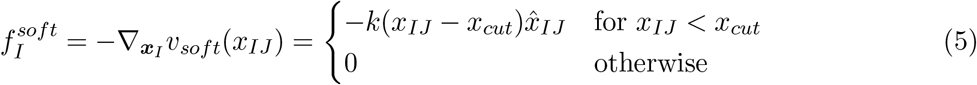
2. One may also encounter cases where a node penetrates a triangulation edge in a time step. In this situation, the resulting tiling may be non-physical. To address this issue, when a cell center *i* is gets very close to a triangulation edge, we artificially increase the viscosity of the cell center as to prevent it from crossing. That is, for a cell center, *I*,

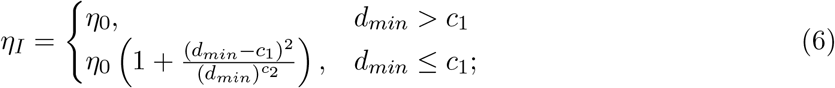

Here *d*_*min*_ is the minimum distance of a cell center *i* from the surrounding triangulation edges. *c*_1_ and *c*_2_ are the regularization parameters. *c*_1_ *>* 0 controls the threshold beyond which the viscosity increases from *η*_0_, and *c*_2_ *>* 0 controls the magnitude of increase. Thus, motion is hindered (i.e., *η*_*i*_ *→ ∞*) only as *d*_*min*_ *→* 0. To determine *c*_1_, we look at the distance between a cell center and its triangulation edge in a given regular hexagonal lattice, which is *l*_0_ sin(*π/*3). Depending on this distance, one can choose *c*_1_ < *l*_0_ sin(*π/*3). We note that this regularization term is only suitable for systems with localized, small-scale cell center motion and causes the system to freeze when attempting long-distance motion. We will relax this constraint and investigate systems with long-range migration in our future work.

### 4 Experimental protocols

#### 4.1 VEGF

HUVECs in passages 3 to 6 were cultured to confluence on fibronectin-coated soft type 1 collagen hydrogels prepared at a concentration of 4 mg/ml as described elsewhere (Antoine et al., 2016). 48 h after seeding, the cells were subjected to vascular endothelial growth factor (VEGF) at a concentration of 50 ng/ml for 66 h. Control experiments consisted of identical protocols in the absence of VEGF. Following VEGF incubation, the cells were fixed with 4% paraformaldehyde and immunostaining performed using standard protocols. Cell-cell junction staining for VE-cadherin was used for cell boundary segmentation. The segmented cell contours were used to determine best-fit ellipses for each cell, and the aspect ratio (AR) was defined as the minor-to-major axis ratio.

#### 4.2 Microgroove substrates

Detailed experimental protocols can be found elsewhere (Leclech et al., 2023). Briefly, substrates with microgrooves of 5 µm width, spacing, and depth were fabricated using photolithography followed by polydimethylsiloxane (PDMS) replication. Flat (no microgrooves) PDMS substrates served as controls. Prior to cell seeding, the PDMS substrates were incubated with fibronectin. Human umbilical vein endothelial cells (HUVECs) in passages 4 to 8 were cultured on the substrates until confluence was reached (typically 24 hours). Cells were then fixed with 4% paraformaldehyde and immunostaining performed using standard protocols. Visualization of cell-cell junctions with VE-cadherin allowed automatic segmentation of cell boundaries. Different cell shape parameters and associated color-coded visual representations were subsequently generated using a custom-made Matlab code. Cell orientation was defined as the mean absolute value of the angle between the major axis of the ellipse that best fits the cell contour and the microgroove direction; 0° corresponds to the microgroove direction and 90° is orthogonal to the microgrooves.

## Footnotes

1

2

